# GC content shapes mRNA decay and storage in human cells

**DOI:** 10.1101/373498

**Authors:** Maïté Courel, Yves Clément, Dominika Foretek, Olivia Vidal Cruchez, Zhou Yi, Marie-Noëlle Benassy, Michel Kress, Caroline Vindry, Marianne Bénard, Clémentine Bossevain, Christophe Antoniewski, Antonin Morillon, Patrick Brest, Arnaud Hubstenberger, Hugues Roest Crollius, Nancy Standart, Dominique Weil

## Abstract

Control of protein expression results from the fine tuning of mRNA synthesis, decay and translation. These processes, which are controlled by a large number of RNA-binding proteins and by localization in RNP granules such as P-bodies, appear often intimately linked although the rules of this interplay are not well understood. In this study, we combined our recent P-body transcriptome with various transcriptomes obtained following silencing of broadly acting mRNA decay and repression factors. This analysis revealed the central role of GC content in mRNA fate, in terms of P-body localization, mRNA translation and mRNA decay. It also rationalized why PBs mRNAs have a strikingly low protein yield. We report too the existence of distinct mRNA decay pathways with preference for AU-rich or GC-rich transcripts. Compared to this impact of the GC content, sequence-specific RBPs and miRNAs appeared to have only modest additional effects on their bulk targets. Altogether, these results lead to an integrated view of post-transcriptional control in human cells where most regulation at the level of translation is dedicated to AU-rich mRNAs, which have a limiting protein yield, whereas regulation at the level of 5’ decay applies to GC-rich mRNAs, whose translation is optimal.

## Introduction

Translation, storage, localization and decay of mRNAs in the cytoplasm are closely coupled processes, which are governed by a large number of RNA-binding proteins (RBPs) [1]. These RBPs have to act in a coordinated manner to give rise to a proteome both coherent with cellular physiology and responsive to new cellular needs. mRNA fate is also intimately linked with their localization in membrane-less organelles, such as P-bodies (PBs). We recently identified the transcriptome and proteome of PBs purified from human cells. Their analysis showed that human PBs are broadly involved in mRNA storage rather than decay [2,3], as also observed using fluorescent decay reporters [4]. However, the mechanism underlying the large but specific targeting of mRNAs to PBs is still unknown, though it clearly results in the co-recruitment of particular RBPs [2].

In mammalian cells, the RNA helicase DDX6, known for its involvement in mRNA decay and translation repression, is a key factor in PB assembly [5]. Human DDX6 interacts with both translational repressors, and the decapping enzyme DCP1/2 and its activators [6,7]. Its yeast homologue Dhh1 is a cofactor of DCP2, as well as a translational repressor [8]. The RBP PAT1B has also been defined as an enhancer of decapping, as it interacts with DDX6, the LSM1-7 heptamer ring and the decapping complex in mammalian cells [9], while in yeast Pat1p activates Dcp2 directly [10] and its deletion results in deadenylated but capped intact mRNA [11,12]. DDX6 and PAT1B interact with the CCR4-NOT deadenylase complex and the DDX6-CNOT1 interaction is required for miRNA silencing [9,13–15]. DDX6 also binds the RBP 4E-T, another key factor in PB assembly, which in turn interacts with the cap-binding factor eIF4E and inhibits translation initiation, including that of miRNA target mRNAs [16]. Altogether, DDX6 and PAT1B have been proposed to link deadenylation/translational repression with decapping. Finally, the 5’-3’ exonuclease XRN1 decays RNAs following decapping by DCP1/2, a step triggered by deadenylation mediated by PAN2/3 and CCR4-NOT or by exosome activity [17].

A number of RBPs also control mRNA fate in a sequence-specific manner, some of them localizing in PBs as well. For instance, the CPEB complex, best described in *Xenopus* oocytes [18], binds the CPE motif in the 3’ untranslated region (UTR) of maternal transcripts through CPEB1, thus controlling their storage and their translational activation upon hormone stimulation [19]. Additional examples include the proteins which bind 3’UTR AU-rich elements (ARE), such as HuR and TTP, to control translation and decay, and play key roles in inflammation, apoptosis and cancer [20]. Protein-binding motifs are generally not unique but are rather defined as consensus sequence elements. In the case of RISC, binding specificity is given by a guide miRNA, which also hybridizes with some flexibility with complementary mRNA sequences. A variety of techniques have therefore been developed to identify the effective RNA targets of such factors, ranging from affinity purification (such as RIP or CLIP) to transcriptome and polysome profiling after RBP silencing, providing the groundwork to address systemic questions about post-transcriptional regulation.

In this study, we addressed the question of the general determinants of mRNA storage and decay in unstressed human cell lines, using the transcriptome of purified PBs and several high-throughput experiments performed after silencing of general storage and decay factors: a polysome profiling after DDX6 silencing, a transcriptome profiling after PAT1B silencing, and two transcriptome profiling after XRN1 silencing. We also used datasets available from the literature, including a transcriptome profiling after DDX6 silencing, a DDX6-CLIP experiment and various datasets reporting RBP and miRNA targets (see Methods). Their combined analysis revealed the central role of mRNA GC content which, by impacting codon usage, PB targeting and RBP binding, influence mRNA fate and contributes to the coordination between two opposite processes: decay and storage.

## Results

### PBs only accumulate AU-rich mRNAs

We have previously shown that PBs store one third of the coding transcriptome in human epithelial HEK293 cells [2]. Such a large transcript number led us to search for general distinctive sequence features that could be involved in PB targeting. As transcript length was reported to be key for mRNA accumulation in stress granules [21], we first analyzed its importance for mRNA accumulation in PBs. When mRNAs were subdivided into six classes ranging from <1.5 kb to >10 kb, longer mRNAs appeared more enriched in PBs than shorter ones, with a clear correlation between length and PB enrichment (Spearman r = 0.39, p-value < 0.0001) (Figures 1A, S1A). However, their increased length in PBs (4.4 kb on average) was less striking than previously observed for stress granule mRNAs (7.1 kb on average) [21].

**Figure 1:**
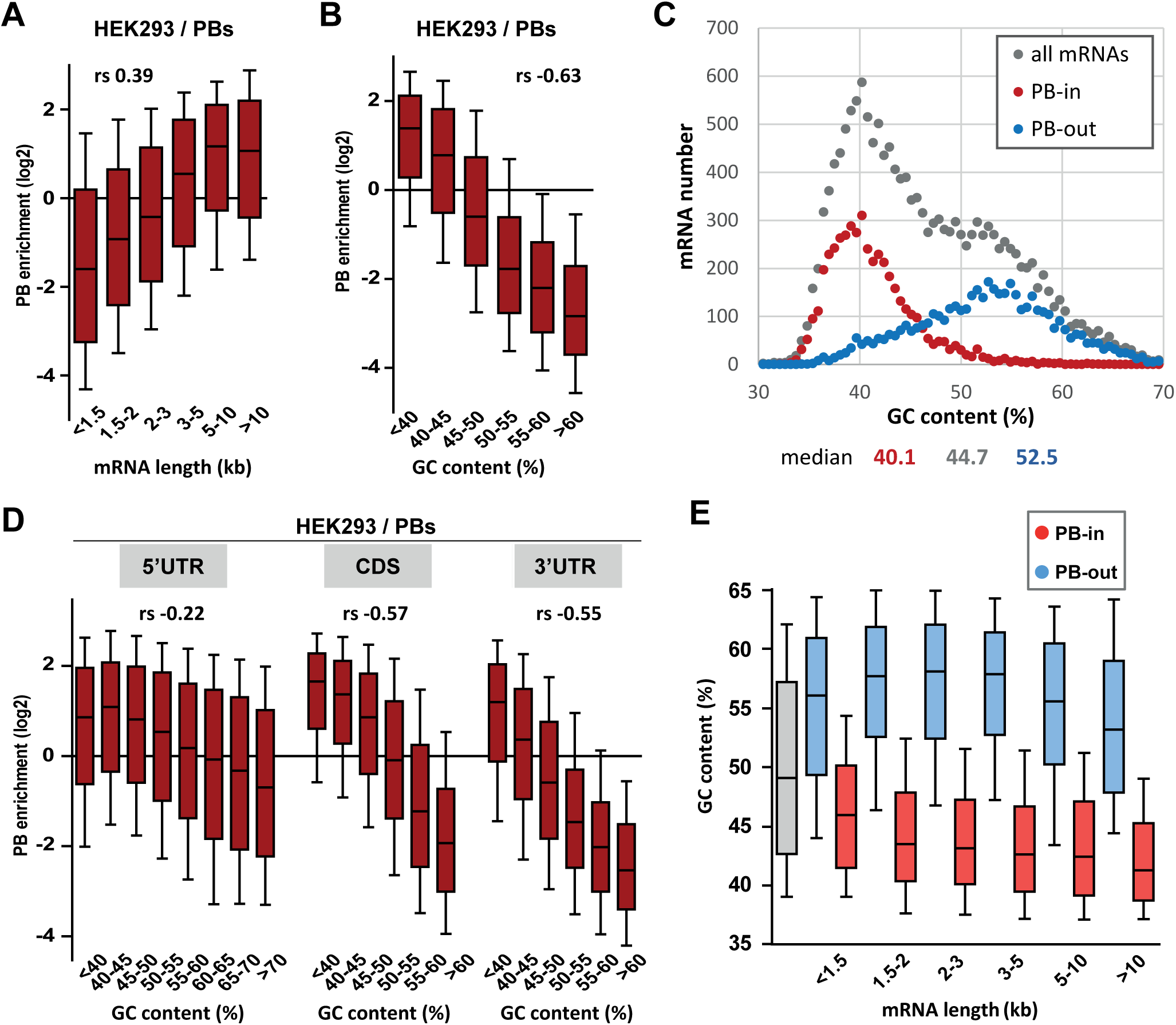
PB mRNAs are AU-rich and longer than average. (A) Long mRNAs are particularly enriched in PBs. Transcripts were subdivided into six classes depending on their length (from <1.5 kb to >10 kb). The boxplots represent the distribution of their respective enrichment in PBs (in dark red). The boxes represent the 25-75 percentiles and the whiskers the 10-90 percentiles. rs is the Spearman correlation coefficient. (B) AU-rich mRNAs are particularly enriched in PBs. Transcripts were subdivided into six classes depending on the GC content of their gene (from <40 to >60%) and analyzed as in (A). (C) PBs only contain transcripts from AU-rich genes. The human transcriptome was subdivided in 77 classes depending on their GC content (0.5% GC increments). The graph represents the number of PB-enriched (PB-in, FC>1, p<0.05, n=4593) and PB-excluded (PB-out, FC<-1, p<0.05, n=4630) transcripts in each class. The distribution of all genes is shown for comparison (in grey, n=15622). The median GC value is indicated below for each group. (D) mRNA localization in PBs mostly depends on the GC content of their CDS and 3’UTR. The analysis was repeated as in (A) using the GC content of the 5’UTR, CDS or 3’UTR, as indicated. For 5’UTRs, the >60% class was subdivided into three classes to take into account their higher GC content compared to CDSs and 3’UTRs. (E) GC content is lower in PB-enriched mRNAs than PB-excluded ones independently of their length. The GC content distribution of PB-enriched (PB-in, p<0.05) and PB-excluded (PB-out, p<0.05) mRNAs was analyzed as in (B). See also Figure S1.

Most remarkably, mRNA accumulation in PBs was strongly dependent on their global nucleotide composition. When genes were subdivided into six classes ranging from <40% to >60% GC, the extent of PB enrichment was predominant for <45% GC and steadily decreased with the GC content (Figures 1B, S1B). While reminiscent of the low GC content reported for stress granule mRNAs in HEK293 cells [21], the correlation was much higher with PB localization (Spearman r = −0.63, p-value < 0.0001) than with stress granule localization (Spearman r = −0.12, p-value < 0.0001) (Figure S1B,C). Indeed, comparing the GC content distribution of the genes whose transcripts are enriched or excluded from PBs with the one of all genes expressed in HEK293 cells, revealed that mRNA storage in PBs is confined to the AU-rich fraction of human genes (Figure 1C).

This raised the possibility that the impact of GC content on PB enrichment resulted indirectly from the genomic context of the genes. To address this issue, we looked at the link between PB enrichment and meiotic recombination, which can influence GC content through GC-biased gene conversion [22]. The correlation between PB enrichment and meiotic recombination was much weaker than between PB enrichment and mRNA GC content (Spearman r = −0.16 vs −0.66, both p-values < 0.0001). Moreover, the latter was almost unchanged when controlling for meiotic recombination (Spearman r = −0.65 vs −0.66, p-value < 0.0001). Finally, it was still significant when controlling for intronic or flanking GC content (Spearman r = −0.33 and −0.45 respectively, all p-values < 0.0001), showing that mRNA base composition and PB enrichment are associated independently of meiotic recombination or the genomic context.

To refine the link between mRNA accumulation in PBs and their GC content, we analyzed separately the influence of their CDS and UTRs. Interestingly, mRNA accumulation in PBs correlated strongly with the GC content of both their CDS and 3’UTR (Spearman r = −0.57 and −0.55, respectively, p-value < 0.0001 for both), and more weakly with the one of their 5’UTR (Spearman r = −0.22, p-value < 0.0001) (Figure 1D, S1B). Moreover, the lower GC content of PB-enriched mRNAs compared to PB-excluded ones was a feature independent of their length, since it was observed in all length ranges (Figure 1E, S1A). Conversely, the higher length of PB mRNAs was a feature independent of their GC content (Figures S1B,D).

In conclusion, while PB mRNAs tend to be longer than average, their most striking feature is that they correspond to an AU-rich subset of the transcriptome.

### GC bias in PBs impacts codon usage and protein yield

The strong GC bias in the CDS of PB mRNAs prompted us to compare the coding properties of PB-stored and PB-excluded mRNAs. Consistently, we found that the frequency of amino acids encoded by GC-rich codons (Ala, Gly, Pro) was lower in PB-stored than in PB-excluded mRNAs, while the frequency of those encoded by AU-rich codons (Lys, Asn) was higher (Figures 2A). The difference could be striking, as illustrated by Lys, whose median frequency in PB-excluded mRNAs was 32% lower than in PB-enriched mRNAs, thus ranging within the lower 17th centile of their distribution (Figures S2A). In addition to the different amino acid composition of encoded proteins, we observed dramatic variation in codon usage between the two mRNA subsets. For all amino acid encoded by synonymous codons, the relative codon usage in PBs versus out of PBs was systematically biased towards AU-rich codons (log2 of the ratio >0, Figure 2B). For example, among the six Leu codons, AAU was used 4-fold more frequently in PB-enriched than in PB-excluded mRNAs, whereas CUG was used 2-fold less frequently. This systematic trend also applied to Stop codons. Some additional codon bias independent of base composition (NNA/U or NNG/C) was also observed for 4 and 6-fold degenerated codons (Figures S2B,C). For instance, Leu was encoded twice more often by CUU than CUA in PB-enriched mRNAs, whereas the use of both codons was low in PB-excluded mRNAs. Similarly, Gly was encoded more often by GGG than GGC in PB mRNAs, whereas the use of both codons was similar in PB-excluded mRNAs (Figure 2C).

**Figure 2:**
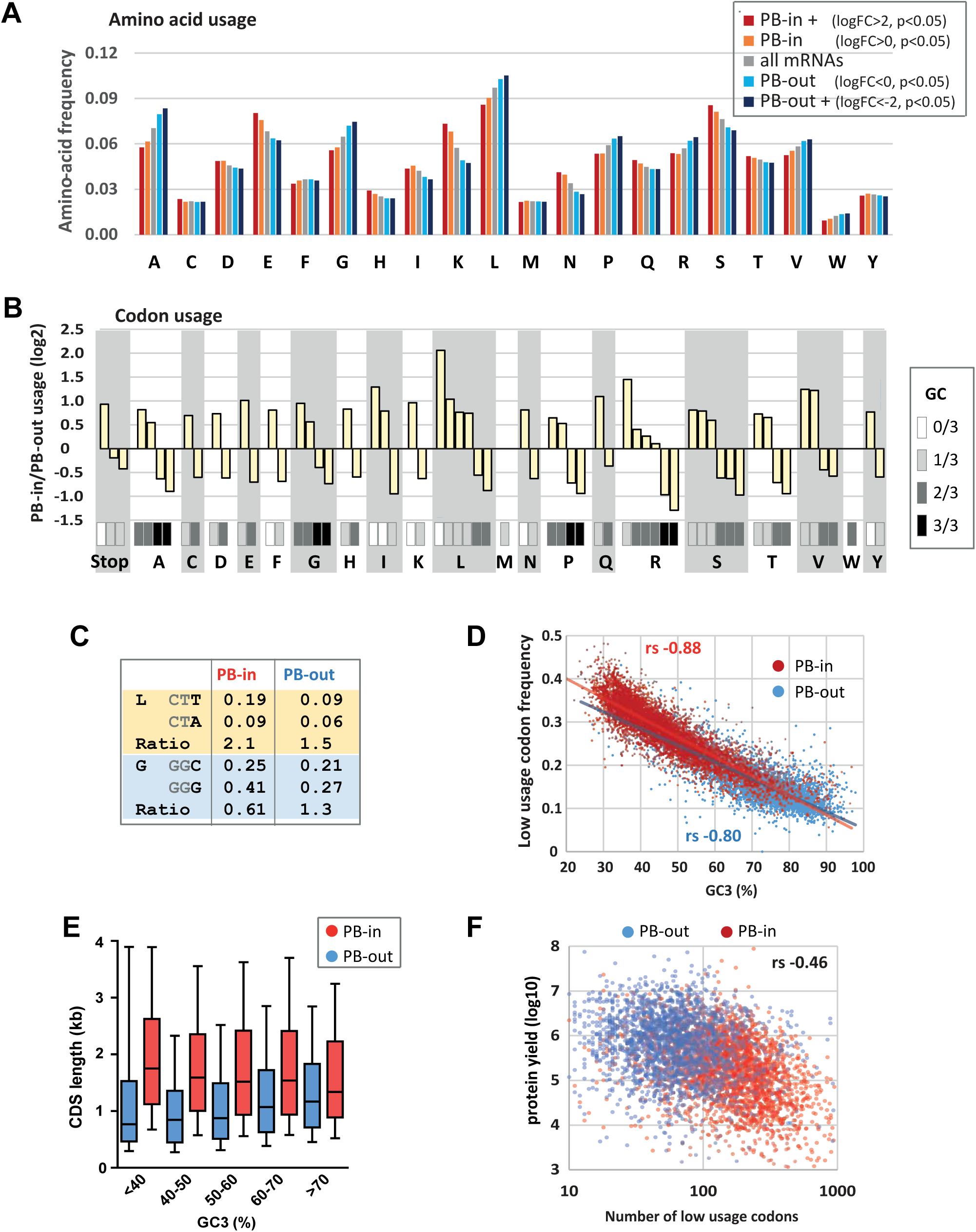
Codon usage is strongly biased in PBs. (A) PB mRNAs and PB-excluded mRNAs encode proteins with different amino acid composition. The graph represents the frequency of each amino acid in the proteins encoded by mRNAs enriched or excluded from PBs, using the indicated PB enrichment thresholds. (B) Codon usage bias in and out of PBs follows their GC content. The relative codon usage for each amino acid was calculated in PB-enriched (PB-in) and PB-excluded (PB-out) mRNAs, using a PB enrichment threshold of +/- 1 (in log2). The graph represents the log2 of their ratio (PB-in/PB-out) and was ranked by increasing values for each amino acid. The GC content of each codon is grey-coded below, using the scale indicated in the right panel. (C) The usage of some codons is biased independently of their GC content. Two examples are shown encoding Leucine (L) and Glycine (G). (D) The frequency of low usage codons is strongly correlated with the GC content of the CDS, independently of their PB localization. The frequency of low usage codons was calculated for mRNAs excluded (PB-out) and enriched (PB-in) in PBs using a PB enrichment threshold of +/- 1 (in log2). It was expressed as a function of the CDS GC content at position 3 (GC3). Note that the slopes of the tendency curves are similar for PB-enriched and PB-excluded transcripts. rs is the Spearman correlation coefficient. (E) PB mRNAs have longer CDS than PB-excluded mRNAs. The analysis was performed as in Figure 1E. (F) The number of low usage codons per CDS is a good determinant of both protein yield and PB localization. The protein yield was expressed as a function of the number of low usage codons for PB-enriched (PB-in) and PB-excluded (PB-out) mRNAs. rs is the Spearman correlation coefficient. See also Figure S2.

In human, the less frequent synonymous codons of an amino acid (normalized relative usage <1, called low usage codons thereafter) generally end with an A or U, with the exception of Thr, Ser, Pro, Ala. Consistent with the GC bias, 23 of the 29 such codons were overrepresented in PB mRNAs (Figure S2D). We calculated the frequency of low usage codons for each CDS, and plotted it as a function of the GC content at the third position (GC3) to avoid any confounding effects of the amino acid bias. As expected, the frequency of low usage codons correlated strongly and negatively with GC3, with AU-rich CDS having a higher frequency of low usage codons than GC-rich CDS (Figure 2D). According to their distinct GC content, PB mRNAs had a higher frequency of low usage codons than PB-excluded mRNAs. However, the correlation coefficient between frequency of low usage codons and GC3 was similar for both mRNA subsets (Spearman r = −0.88 for PB-enriched; −0.80 for PB-excluded mRNAs, p-value < 0.0001 for both), meaning that their different frequency of low usage codons could be largely explained by their GC bias alone.

We previously reported that protein yield, defined as the ratio between protein and mRNA abundance in HEK293 cells, was 20-times lower for PB-enriched than PB-excluded mRNAs. This was not due to translational repression within PBs, as the proportion of a given mRNA in PBs hardly exceeded 15%, but rather to some intrinsic mRNA property [2]. In this respect, the frequency of low usage codons correlated well with PB localization (Spearman r = 0.59, p-value < 0.0001) but moderately with protein yield (Spearman r = −0.21, p-value < 0.0001) (Figure S2E). Conversely, the CDS length correlated well with protein yield (Spearman r = −0.43, p-value < 0.0001) and moderately with PB localization (Spearman r = 0.26, p-value < 0.0001) though nevertheless independently of their GC3 content (Figure 2E). Finally, combining the frequency of low usage codons with the CDS length, that is, considering the absolute number of low usage codons per CDS, was a shared parameter of both protein yield (Spearman r = −0.46, p-value < 0.0001, Figure 2F) and PB localization (Spearman r = 0.49, p-value < 0.0001). Strikingly, CDS with more than 100 low usage codons were particularly enriched in PBs, while those under 100 were mostly excluded (Figure 2F).

In conclusion, the strong GC bias in PB mRNAs results in both a biased amino acid composition of encoded proteins and a biased codon usage. Furthermore, the high number of low usage codons in PB mRNAs is a likely determinant of their low protein yield.

### The PB assembly factor DDX6 has opposite effects on mRNA decay and translation depending on their GC content

In human, the DDX6 RNA helicase is key for PB assembly [5]. It associates with a variety of proteins involved in mRNA translation repression and decapping [6,7], suggesting that it plays a role in both processes. To investigate how DDX6 activity is affected by mRNA GC content, we conducted a polysome profiling experiment in HEK293 cells transfected with DDX6 or control β-globin siRNAs for 48h. In these conditions, DDX6 expression decreased by 90% compared to control cells (Figure S3A). The polysome profile was largely unaffected by DDX6 silencing, implying that DDX6 depletion did not grossly disturb global translation (Figure S3B). Polysomal RNA isolated from the sucrose gradient fractions (Figure S3B) and total RNA were used to generate libraries using random hexamers to allow their amplification independently of the length of their poly(A) tail. As expected, both total and polysomal DDX6 mRNA was markedly decreased (by 72%) (Figure S3C-E; Table S1, sheet1). Since DDX6 is cytoplasmic [23], we assumed that total mRNA accumulation generally reflected their increased stability. As polysomal accumulation can result from both regulated translation and a change in total RNA without altered translation, we then used the polysomal to total mRNA ratio as a proxy measurement of translation rate.

Analysis of the whole transcriptome showed a link between mRNA fate following DDX6 depletion and their GC content, but, intriguingly, the correlation was positive for changes in total RNA (Spearman r = 0.44, p-value < 0.0001; Figure S3F) and negative for changes in polysomal RNA (Spearman r = −0.27, p-value < 0.0001; Figure S3G). Therefore, DDX6 depletion affected different mRNA subsets in total and polysomal RNA. The extent of mRNA stabilization steadily increased with the GC content and became predominant for transcripts with >50% GC (Figures 3A, left panel, and S4A). This analysis was repeated on an independent dataset available from the ENCODE project [24], obtained in a human erythroid cell line, K562, following induction of a stably transfected DDX6 shRNA, and using an oligo(dT)-primed library. Despite the differences in cell type, depletion procedure and sequencing methods, again, mRNA stabilization preferentially concerned those with high GC content (Spearman r = 0.59, p-value < 0.0001; Figures 3A, right panel, and S4A; Table S1, sheet2). In contrast, following DDX6 silencing in HEK293 cells, the translation rate predominantly increased for transcripts with less than 45% GC (Spearman r = −0.53, p-value < 0.0001; Figure 3B). As a result, mRNAs with the most upregulated translation rate were the least stabilized, and conversely (Figure S4B).

**Figure 3:**
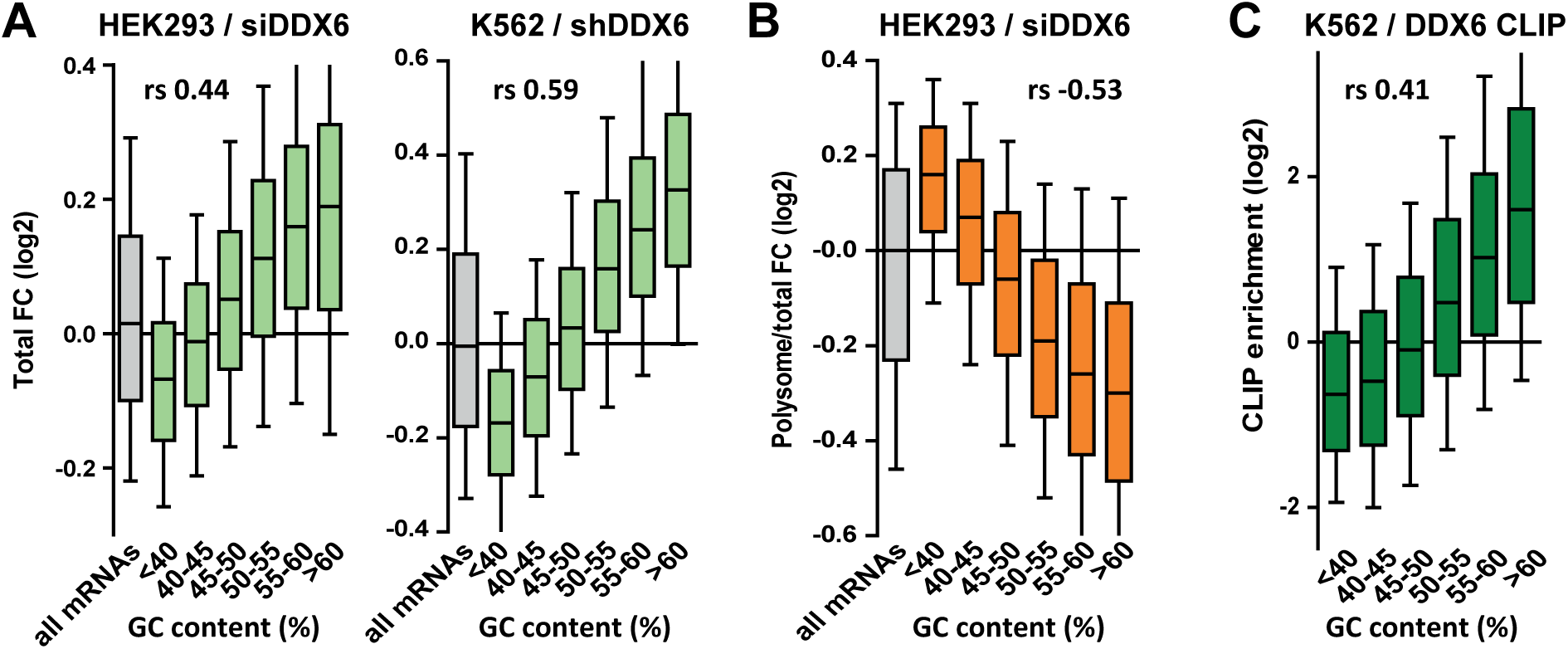
DDX6 silencing has opposite effects on mRNA decay and translation depending on the mRNA GC content. (A) mRNA stabilization after DDX6 silencing in HEK293 and K562 cells applies to GC-rich mRNAs. The fold-changes (FC) in mRNA accumulation (in green) were analyzed as in Figure 1B. (B) mRNA translation derepression after DDX6 silencing in HEK293 cells applies to AU-rich mRNAs. The fold-changes in translation rate (in orange) were analyzed as in (A). (C) GC-rich mRNAs are particularly enriched in the DDX6 CLIP experiment (in dark green). See also Figures S3-5.

To investigate how DDX6 activity was related to its binding to RNA, we used the CLIP dataset of K562 cells, also available from the ENCODE project. In both HEK293 and K562 cells, the mRNAs clipped to DDX6 were particularly stabilized after DDX6 knockdown, as compared to all mRNAs (Figure S4C; Table S1, sheet3), while they were not regulated at the level of translation in HEK293 cells (Figure S4D). In agreement, mRNAs with a high GC content were strongly enriched in the DDX6 CLIP experiment (Spearman r = 0.41, p-value < 0.0001; Figures 3C and S4A). Then, as we previously showed that DDX6 can oligomerize along repressed transcripts [25], we also considered mRNA length. While DDX6-dependent decay had a marginal preference for short transcripts (Spearman r = −0.09, p < 0.0001), as a combined effect of CDS and 3’UTR length (Figure S4E and S4F), DDX6-dependent translation repression was independent of the CDS length but higher on mRNAs with long 3’UTRs (Spearman r = 0.16, p < 0.0001; Figure S4E and S4G). Interestingly, the GC content of the CDS and the 3’UTR were similarly predictive of DDX6 sensitivity, whether for mRNA decay (Spearman r = 0.42 and 0.40 for CDS and 3’UTR, respectively, p < 0.0001 for both) or for translation repression (Spearman r = −0.53 and −0.52, respectively, p < 0.0001 for both), while the 5’UTR was less significant (Spearman r = 0.18 and −0.15 for decay and translation repression, respectively, p < 0.0001 for both; Figures S5A-C).

Altogether, we showed that DDX6 knockdown affected differentially the mRNAs depending on the GC content of both their CDS and 3’UTR, with the most GC-rich mRNAs being preferentially regulated at the level of decay and the most AU-rich mRNAs at the level of translation.

### DDX6/XRN1 and PAT1B enhance the decay of separate sub-classes of mRNAs with distinct GC content

DDX6 acts as an enhancer of decapping to stimulate mRNA decay, upstream of RNA degradation by the XRN1 5’ to 3’ exonuclease. To investigate whether XRN1 targets are similarly GC-rich, we performed XRN1 silencing experiments in two cell lines. HeLa cells were transfected with XRN1 siRNA (Figure S6A; Table S1, sheet4), while HCT116 cells stably transfected with an inducible XRN1 shRNA were induced with doxycyclin (Figure S6B; Table S1, sheet5), both for 48h. In both cell lines, XRN1-dependent decay preferentially acted on mRNAs which were GC-rich (Spearman r = 0.41 for HeLa and 0.49 for HCT116, p < 0.0001 for both; Figures 4A and S4A) and localized out of PBs (Spearman r = - 0.35, p<0.0001; Figure S6C), as observed for DDX6.

**Figure 4:**
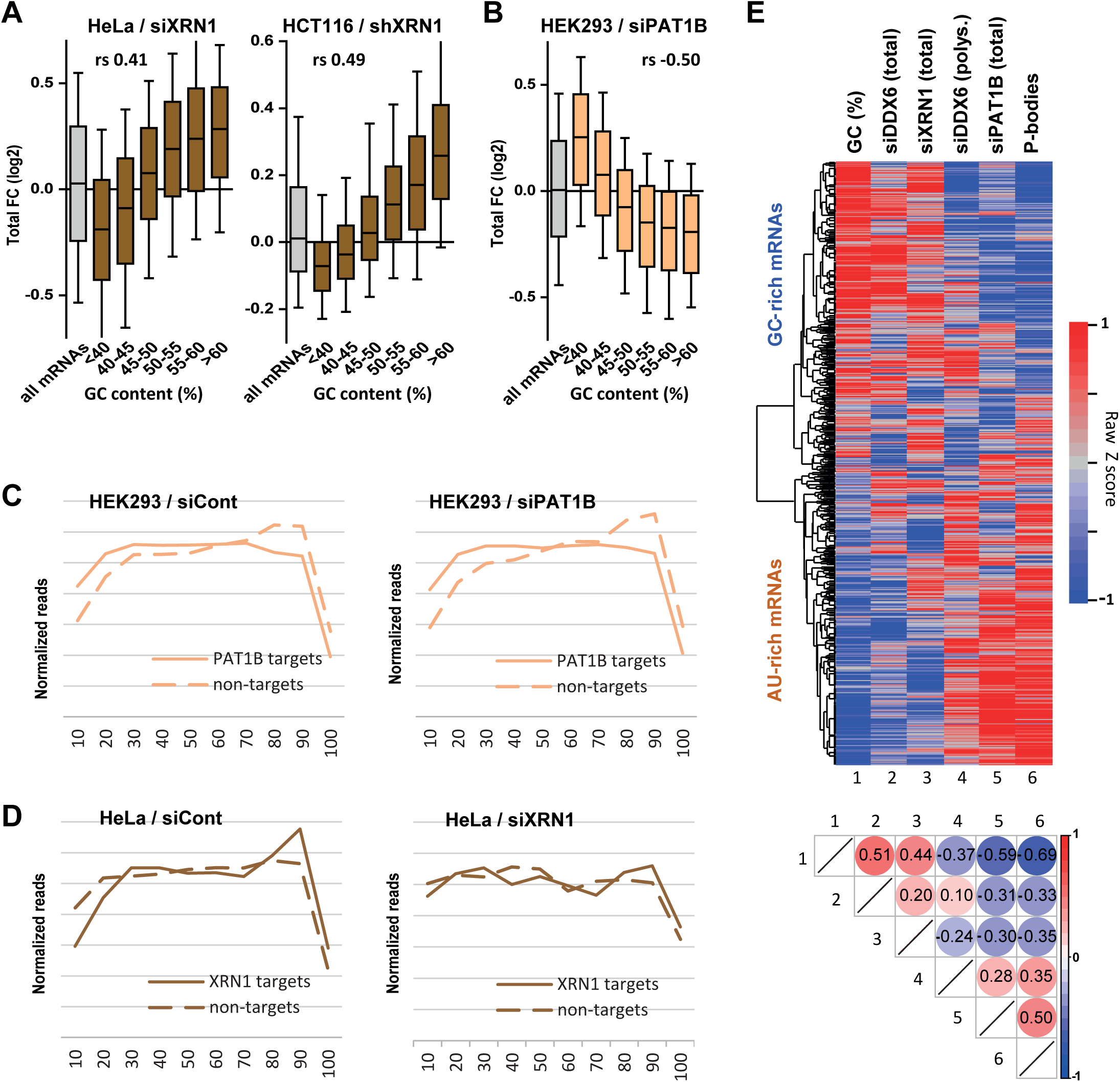
XRN1 and PAT1B involvement in mRNA decay depends on their GC content. (A) mRNA stabilization after XRN1 silencing in HeLa and HCT116 cells (in brown) applies to GC-rich mRNAs. The analysis was performed as in Figure 1B. The GC content distribution for all mRNAs is presented for comparison (in grey). (B) mRNA stabilization after PAT1B silencing in HEK293 cells (in peach) applies to AU-rich mRNAs. The analysis was performed as in (A). (C) Read coverage of XRN1 targets (FC>1, n=333) and non-targets (FC<-1, n=139), as defined in the siXRN1 dataset. Their average read coverage was analyzed in control cells (upper panel) and after XRN1 silencing (lower panel), and normalized as described in the Methods. (D) Read coverage of PAT1B targets (FC>0.6, n=616) and non-targets (FC<-0.6, n=493), as defined in the siPAT1B dataset. The data were analyzed as in (C). (E,F) Clustering analysis of mRNAs depending on their GC content, their differential expression after silencing DDX6, XRN1 or PAT1B, and their enrichment in PBs. Raw GC content and log2 transformed ratio of the other datasets were used for the clustering of both transcripts (lines) and datasets (columns). The values were color-coded as indicated on the right scale, and the Spearman correlation matrix is presented in F (all p-values <10-48). The heatmap highlights the distinct fate of GC-rich and AU-rich mRNAs. See also Figures S4 and S6.

PAT1B is a well-characterized direct DDX6 partner known for its involvement in mRNA decay [9,26– 28]. However, using our previous PAT1B silencing experiment in HEK293 cells [9], we surprisingly found a negative correlation between mRNA stabilization after PAT1B and after DDX6 silencing (Spearman r = −0.31, p<0.0001; Figure S6D; Table S1, sheet6), suggesting that they largely target separate sets of mRNAs. Unexpectedly, the correlation was however positive with translational derepression after siDDX6 (Spearman r = 0.45, p<0.0001; Figure S6E), indicating that PAT1B preferentially targets mRNAs that are translationally repressed by DDX6. Accordingly, these transcripts are prone to PB storage (Spearman r = 0.49, p<0.0001; Figure S6F), as reported previously [9]. Indeed, in contrast to DDX6 and XRN1 decay targets, PAT1B targets were strikingly AU-rich (Spearman r = −0.50, p < 0.0001; Figures 4B and S4A). To gain insight into the mechanism of PAT1B-dependent decay, we analyzed the read coverage of PAT1B targets. Prior to any silencing, these transcripts had a higher 5’ coverage than non-target mRNAs (Figure 4C, left panel), and it remained high after PAT1B silencing (Figure 4C, right panel). In contrast, XRN1 targets had a lower 5’ coverage than non-targets prior to silencing (Figure 4D, left panel), and this difference of coverage was suppressed following XRN1 depletion, as expected from a 5’ decay pathway (Figure 4D, right panel). These results suggest that mRNA stabilization in the absence of PAT1B does not result from their 5’ end protection.

In conclusion, DDX6 and PAT1B enhance the decay of distinct mRNA subsets, which strongly differ in their GC content. The results suggest that DDX6 is a cofactor of XRN1 5’ 3’ exonuclease, whereas PAT1B-dependent decay likely relies on 3’ to 5’ degradation.

To obtain a global visualization of the results we conducted a clustering analysis of the various datasets (Figure 4E). Note that to avoid clustering interdependent datasets, we included the changes in polysomal RNA after DDX6 silencing rather than in translation rate. Altogether, the heatmap shows that GC-rich mRNAs are excluded from PBs and tend to be decayed by a mechanism involving DDX6 and XRN1, while AU-rich mRNAs are recruited in PBs, they undergo DDX6-dependent translation repression and their decay depends on PAT1B.

### Specific mRNA decay factors and translation regulators target mRNAs with distinct GC content

Having shown that GC content is a distinctive feature of DDX6- and XRN1-versus PAT1B-dependent mRNA decay, we investigated the link between this global sequence determinant and a variety of sequence-specific post-transcriptional regulators for which relevant genome-wide datasets are available (Figure S7A).

On the mRNA decay side (group I lists), we considered the Nonsense Mediated Decay (NMD) pathway, taking as targets the mRNAs cleaved by SMG6 [29], and the m6A-associated decay pathways, using the targets of the YTHDF2 reader defined by CLIP [30,31]. We also analyzed mRNAs with a 5’UTR-located G4 motif, which have been shown to be preferential substrates of murine XRN1 in vitro [32]. On the translation regulation side (group II lists), we analyzed the TOP mRNAs, whose translation is controlled by a TOP motif at the 5’ extremity [33], and targets of various PB proteins and/or DDX6 partners: FXR1-2, FMR1, PUM1-2, IGF2BP1-3, the helicase MOV10, ATXN2, 4E-T, ARE-containing mRNAs and the targets of the two ARE-binding proteins HuR and TTP [2,6]. We also included mRNAs with a CPE motif, since DDX6 is a component of the CPEB complex that binds CPEs [18]. Of note, among the group II factors, some are known to also affect mRNA half-life, as exemplified by the ARE-binding proteins [20]. G4, ARE and CPE motifs have been defined in silico, while the targets of the various factors originate from RIP and CLIP approaches or mouse studies in the case of TOP mRNAs (see Methods).

Intriguingly, compared to all mRNAs, group I list mRNAs were GC-rich, as well as TOP mRNAs and ATXN2 targets, whereas all other group II lists were AU-rich (Figure 5A). Furthermore, they shared common behavior in the various experiments. This is summarized in Figure 5B in a heatmap representing their median value in each dataset, while Figures S7 and S8 provide detailed analysis, as described below.

**Figure 5:**
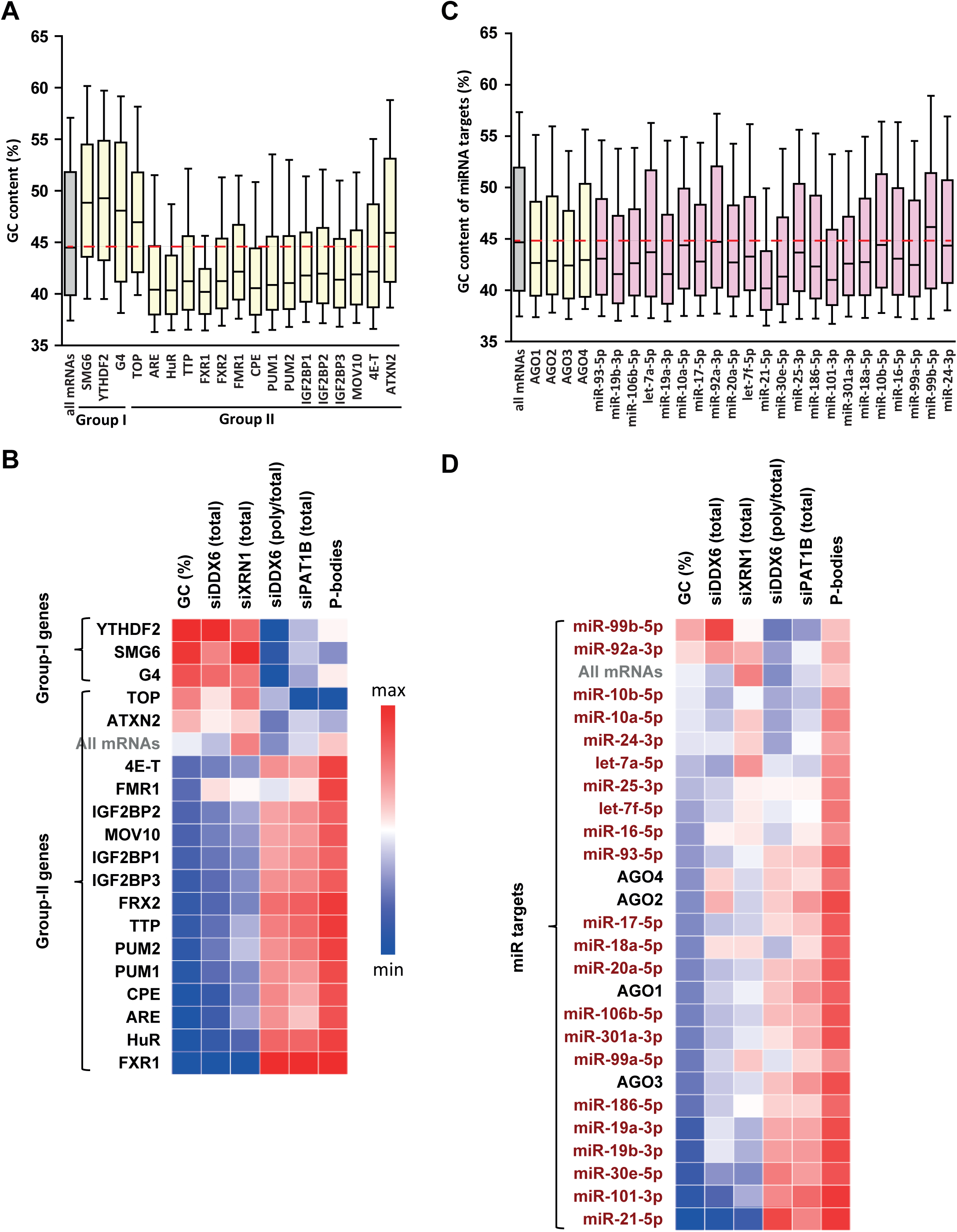
GC biases in the targets of various RNA decay factors and translation regulators. (A) GC content biases in the targets of various RBPs. The targets of the indicated factors were defined using CLIP experiments or motif analysis (see methods). The boxplots represent the distribution of the GC content of their gene (in yellow). The distribution for all mRNAs is presented for comparison (in grey) and the red dashed line indicates its median value. (B) Heatmap representation of the different factors depending on the behavior of their mRNA targets in the different datasets. The lines were ordered by increasing GC content, and the columns as in Figure 4E. (C) GC content biases in the targets of various AGO proteins and miRNAs. The AGO targets (in yellow) were defined using CLIP experiments and the miRNAs targets (in violet) using miRTarbase. The data were represented as in (A). (D) Heatmap representation of AGO and miRNAs depending on the behavior of their mRNA targets in the different datasets. The data were represented as in (B), using the same color code. See also Figures S7-11.

Group I list mRNAs tended to be dependent on DDX6 and XRN1 but not on PAT1B for decay (Figure S7B-D), with nevertheless some variation between cell lines, as only SMG6 targets were sensitive to XRN1 depletion in HeLa cells (Figure S7C, upper panel). They did not accumulate in PBs and their translation was independent of DDX6 (Figure S8A,B). These results were consistent with their high GC content and our global analysis above. However, surprisingly, within PB-excluded mRNAs, there was little or no additional effect of being a SMG6 target, an YTHDF2 target or containing a G4 motif, neither for DDX6-nor for XRN1-dependent decay (Figures S8C,D).

Group II list mRNAs, except TOP mRNAs, ATXN2 and 4E-T targets, had the exact mirror fate compared to group I lists: they were stabilized following PAT1B silencing (Figure S7D), as previously reported for ARE-containing mRNAs and the targets of the ARE-BPs HuR and TTP [9], but not following DDX6 or XRN1 silencing (Figure S7B,C); they were enriched in PBs and translationally more active after DDX6 silencing (Figure S8A,B), which is consistent with the reported presence of most of these regulatory proteins in PBs [2,34].

Among the three group II outsiders, ATXN2 targets and TOP mRNAs behaved like group I lists, except that they were not dependent on DDX6 for decay. ATXN2 is a major DDX6 partner excluded from PBs [6,35], which is consistent with its targets being also excluded from PBs (Figure S8A). However, these mRNAs were weakly or not stabilized following DDX6 or XRN1 silencing (Figure S7B,C), leaving unresolved the function of the ATXN2/DDX6 interaction in the cytosol. TOP mRNAs are special within group II lists, in that their translational control relies on a cap-adjacent motif rather than on 3’UTR binding. 4E-T is a major DDX6 partner required for PB assembly [6,16]. Intriguingly, its targets were markedly enriched in PBs (Figure S8A), though poorly affected by PAT1B silencing in terms of stability (Figure S7D), or by DDX6 silencing in terms of translation (Figure S8B). This dissociation between PB localization and mRNA fate indicated that PB recruitment is not sufficient to determine PAT1B-dependent decay and DDX6-dependent translation repression. It also pointed to a particular role of 4E-T in PB targeting or scaffolding.

Our analysis is also informative on the link between DDX6-dependent decay and codon usage. Previous yeast studies have debated whether suboptimal codons could enhance DDX6 recruitment to trigger mRNA decay [36] or not [37,38]. We showed above that GC-rich mRNAs, which tend to be decayed by DDX6 and to bind DDX6 (Figure 3A,C), are enriched for optimal rather than suboptimal codons (Figure 2D). Furthermore, the correlation between polysomal retention (defined as the fraction of total mRNA present in polysomes) and DDX6-dependent decay was weak (Spearman r = 0.10, p<0.0001; Figure S8E). Thus, in HEK293 cells, this mechanism seemed to account for a minor part of DDX6-dependent decay, if any, as in mouse stem cells [39].

In conclusion, we observed that mRNA decay regulators preferentially target GC-rich mRNAs and trigger DDX6- and XRN1-dependant decay, whereas most translation regulators preferentially target AU-rich mRNAs, resulting in storage in PBs and sensitivity to PAT1B-dependent decay.

### Targets of the miRNA pathway have a biased GC content

The miRNA pathway, which leads to translation repression and mRNA decay, has been previously associated with DDX6 activity and PB localization [40,41]. To study this pathway, we used the list of AGO1-4 targets, as identified in CLIP experiments [31] (Figure S9A). In addition, we analyzed the experimentally documented targets of the 22 most abundant miRNAs in HEK293 cells (19 from [42], and 3 additional ones from our own quantitation, Figure S9B), as described in miRTarBase [43]. The mRNA targets of AGO proteins were AU-rich, as observed for most group II RBPs, and this was also true for the targets of most miRNAs when analyzed separately (Figure 5C). Overall, they also shared common behavior in the various silencing experiments and PB dataset, with nevertheless some differences. This is summarized in Figure 5D in a heatmap representing their median value in each dataset, while Figure S9 provides detailed analysis, as described below.

The AGO targets tended to accumulate in PBs (Figure S9C; note that the number of AGO4 targets was too small to reach statistical significance) and their translation was DDX6-dependent (Figure S9D). In terms of stability, only AGO2 targets were marginally DDX6-dependent (Figure S9E), and the effects were not stronger when analyzing separately the mRNAs enriched or excluded from PBs (Figure S9F). In contrast, their decay was PAT1B-dependent (Figure S9G).

The targets of the 22 miRNAs had a behavior overall similar to the targets of AGO proteins, with accumulation in PBs, DDX6-dependent translation, and PAT1B rather than DDX6- or XRN1-dependent stability (Figure 5D). However, our analysis revealed some differences between miRNAs, particularly in terms of extent of PB storage (Figure S9H), which appeared associated with distinct GC content: at the two extremes, miR21-5p targets were particularly AU-rich and strongly enriched in PBs, while the targets of miR-99b-5p, the most GC-rich in these sets, were not. In terms of translation, miR-18a-5p targets were not sensitive to DDX6 silencing, despite clear enrichment in PBs. In terms of stability, the targets of the less GC-rich miR-99b-5p were sensitive to DDX6 but not PAT1B silencing.

In conclusion, miRNA targets generally tend to be AU-rich, like the targets of most translation regulators, and accumulate in PBs. Their stability is not markedly affected following DDX6 or XRN1 silencing, but is dependent on PAT1B.

### The GC content of mRNAs shapes post-transcriptional regulation

As the global GC content appeared closely linked to mRNA fate, but also to RBP and miRNA binding, as well as to translation activity, we then conducted analysis aiming at ranking the importance of these various features.

We first assessed the respective weight of the GC content and the binding capacity of particular RBPs. To this aim, we ordered the whole transcriptome according to their GC content before subdivision in groups of 500 transcripts, and we plotted the median fold-change of each group in our various RNAseq datasets as a function of its median GC content. The same values were then calculated for the various group I and II target lists and overlaid for comparison (Figure S10). Surprisingly, we observed that being targets of particular RBPs generally did not markedly modify the behavior forecasted from their GC content. This was particularly true for DDX6- and XRN1-dependent decay, with only 4E-T targets being more stabilized after DDX6 silencing than expected from their GC content (Figures S10A,B). In terms of translation, only FXR1 targets were slightly more translated after DDX6 depletion than expected from their GC content (Figure S10C). They were also more dependent on PAT1B for decay (Figure S10D) and more enriched in PBs (Figure S10E). In the case of HuR, TTP, FXR1-2, FMR1, PUM2, IGF2BP1-3 and MOV10, there was some, but minimal, additional PAT1B-sensitivity and PB enrichment.

Similarly, the fate of the miRNA targets was mostly in the range expected from their GC content (Figure S11). Nevertheless, some miRNA-specific effects were observed. For instance, the targets of several miRNAs were more stabilized than expected after DDX6 depletion, including miR-99b-5p, 92a-3p, 16-5p, 18a-5p, 19a-3p, 19b-3p (Figure S11A), though this was not observed following XRN1 depletion (Figure S11B). Similarly, while the targets of miR-101-3p and miR-21-5p were both particularly enriched in PBs (Figure S11E), only miR-101-3p targets were particularly dependent on PAT1B for decay (Figure S11D). Interestingly, we noted that the median GC content of the miRNA targets correlated with the GC content of the miRNA itself (Figure S11F). Thus, despite their small size, the miRNA binding sites tend to have a GC content similar to that of their full-length host mRNA, which affects their fate in terms of PB localization and post-transcriptional control.

Altogether, our analysis showed that the dominant parameter in steady-state conditions appeared to be the mRNA GC content, with the regulatory proteins making subsidiary contributions. The high correlation observed between GC content and PB localization raised the possibility that localization out of PBs was sufficient to determine mRNA sensitivity to DDX6 and XRN1 decay. While a tempting hypothesis, it could not explain all DDX6- and XRN1-dependent decay, since TOP mRNAs were strongly excluded from PBs (Figure S8A), but unaffected by DDX6 or XRN1 silencing (Figure S7B,C). To address this issue more generally, we considered the fate of the minor subset of AU-rich mRNAs that were excluded from PBs. Compared to other similarly AU-rich transcripts, these mRNAs were indeed more sensitive to XRN1-dependent decay (Figure S12A). However they were not more sensitive to DDX6-dependent decay (Figure S12B). This suggested that XRN1 preference for GC-rich mRNAs is at least in part related to their exclusion from PBs, whereas DDX6 has a true preference for GC-rich mRNAs.

Another major issue was to distinguish the contribution of CDS and 3’UTR to PB localization, since they have very similar GC contents (Spearman r=0.72, p<0. 0001). Indeed, PB targeting could be driven by the CDS GC content, as a feature important for codon usage, and hence translation efficiency. Alternatively, it could be driven by the 3’UTR GC content, as a feature important for the binding of particular RBPs. As a first approach, we analyzed PB localization of long non-coding RNAs (lncRNAs) [2]. The correlation between their GC content and PB accumulation was significant (Spearman r=-0.20, p<0.0001), but much weaker than observed for mRNAs (−0.66) or 3’UTRs (−0.55). In fact, AU-rich lncRNAs weakly accumulated in PBs, while GC-rich lncRNAs were markedly excluded (Figure S12C,D). As a second approach, we directly analyzed the respective contribution of the GC content of CDS and 3’UTR to PB localization. On one side, we analyzed transcripts by groups of similar 3’UTR GC content. Their GC3 was systematically much lower in PB mRNAs than in PB-excluded mRNAs, with differences ranging from 9 to 13% GC (Figures 6A and S12D). In a mirror analysis, we analyzed groups of transcripts with similar GC3. The importance of the 3’UTR GC content became visible only for GC3 higher than 50% GC (note that GC3 median value is 59% GC), with AU-rich 3’UTR allowing for their accumulation in PBs despite a GC-rich CDS (Figures 6B and S12D). We concluded that PB localization is driven by both the CDS and the 3’UTR GC content, with the CDS being the primary determinant.

**Figure 6:**
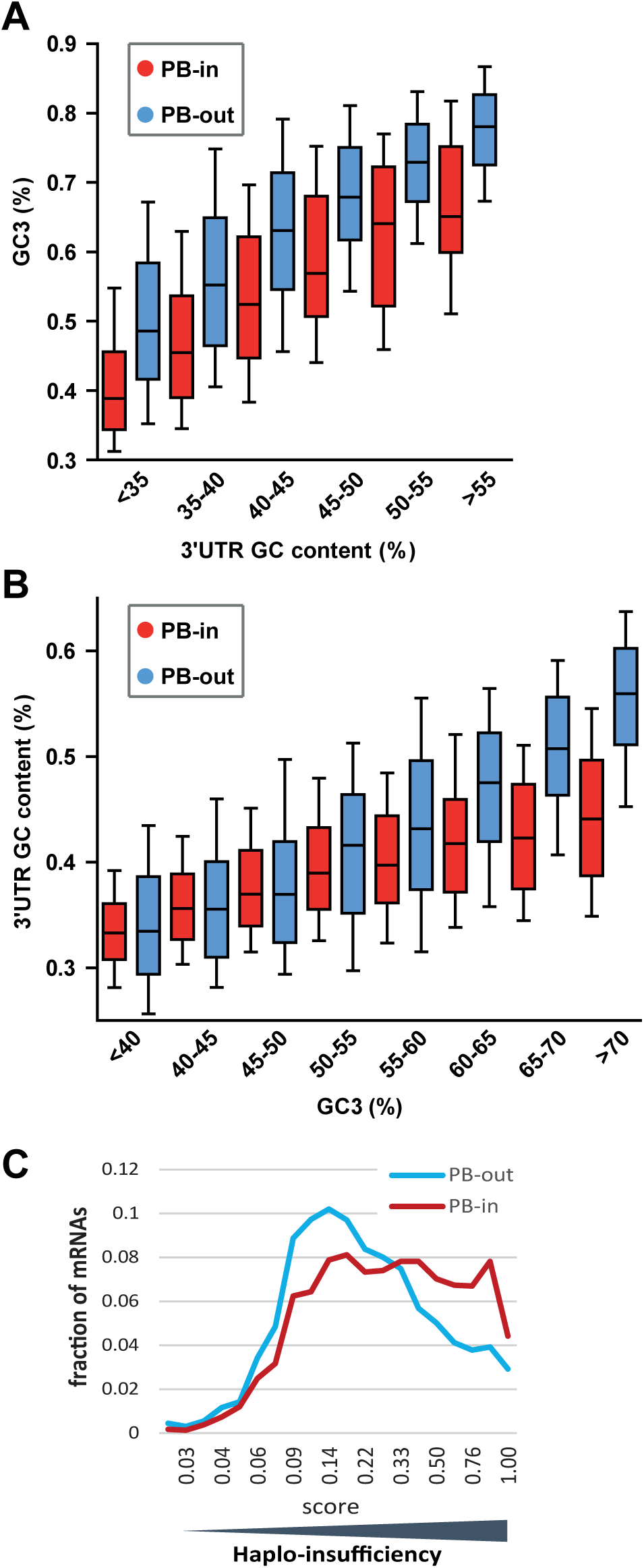
The GC content of the CDS and the 3’UTR are the primary and secondary determinants of PB localization, respectively. (A) General importance of the CDS. Transcripts were subdivided into six classes depending on the GC content of their 3’UTR (from <40 to >55%). The boxplots represent the distribution of the GC content of their CDS at position 3 (GC3) in PB-enriched (PB-in) and PB-excluded (PB-out) mRNAs. (B) Importance of the 3’UTR for GC-rich CDS. Transcripts were subdivided into eight classes depending on their GC3 (from <40 to >70%). The boxplots represent the distribution of the GC content of their 3’UTR in PB-enriched (PB-in) and PB-excluded (PB-out) mRNAs. (C) The transcripts of haplo-insufficiency genes are enriched in PBs. The haplo-insufficiency score is the probability that a gene is haplo-insufficient, as taken from the Huang et al. study [44]. The analysis was performed for PB-enriched (PB-in, n=4646) and PB-excluded (PB-out, n=4205) mRNAs. The results were similar using Steinberg et al. scores [45]. See also Figure S12.

Altogether, our results pointed to independent contributions of both CDS and 3’UTR regions to PB localization, CDS likely acting through codon usage and low translation efficiency, and 3’UTR through the binding to RBPs with affinity for AU-rich sequences. We speculate that suboptimal translation makes mRNAs optimal targets for translation regulation, since any control mechanism has to rely on a limiting step. Conversely, optimally translated transcripts would be better controlled by decay. One prediction of this model is that proteins produced in limiting amounts, such as those encoded by haplo-insufficiency genes, are more likely to be encoded by PB mRNAs. Genome-wide haplo-insufficiency prediction scores have been defined for human genes, using diverse genomic, evolutionary, and functional properties trained on known haplo-insufficient and haplo-sufficient genes [44,45]. Using these scores, we found that haplo-insufficient mRNAs were indeed enriched in PBs (Figure 6C).

## Discussion

### An integrated model of post-transcriptional regulation

Our combined analysis of the transcriptome of purified PBs together with transcriptomes following the silencing of broadly-acting storage and decay factors, including DDX6, XRN1 and PAT1B, provided a general landscape of post-transcriptional regulation in human cells, where mRNA GC content plays a central role. As schematized in Figure 7, GC-rich mRNAs are excluded from PBs and mostly controlled at the level of decay, by a mechanism involving the helicase DDX6 and the 5’ exonuclease XRN1. In contrast, AU-rich mRNAs are enriched in PBs and mostly controlled at the level of translation, by a mechanism also involving DDX6, and at the level of decay by the DDX6 partner PAT1B, a decay likely to be from the 3’ extremity. Accordingly, NMD and m6A-associated mRNA decay pathways tend to target GC-rich mRNAs, while most sequence-specific translation regulators and miRNAs tend to target AU-rich mRNAs. The distinct fate of GC-rich and AU-rich mRNAs correlates with a contrasting protein yield resulting from both different codon usage and CDS length. Thus, 5’ mRNA decay appeared to be used to control preferentially mRNAs with optimal translation, which are mostly GC-rich, whereas translation regulation is mostly used to control mRNAs with limiting translational efficiency, which are AU-rich.

**Figure 7:**
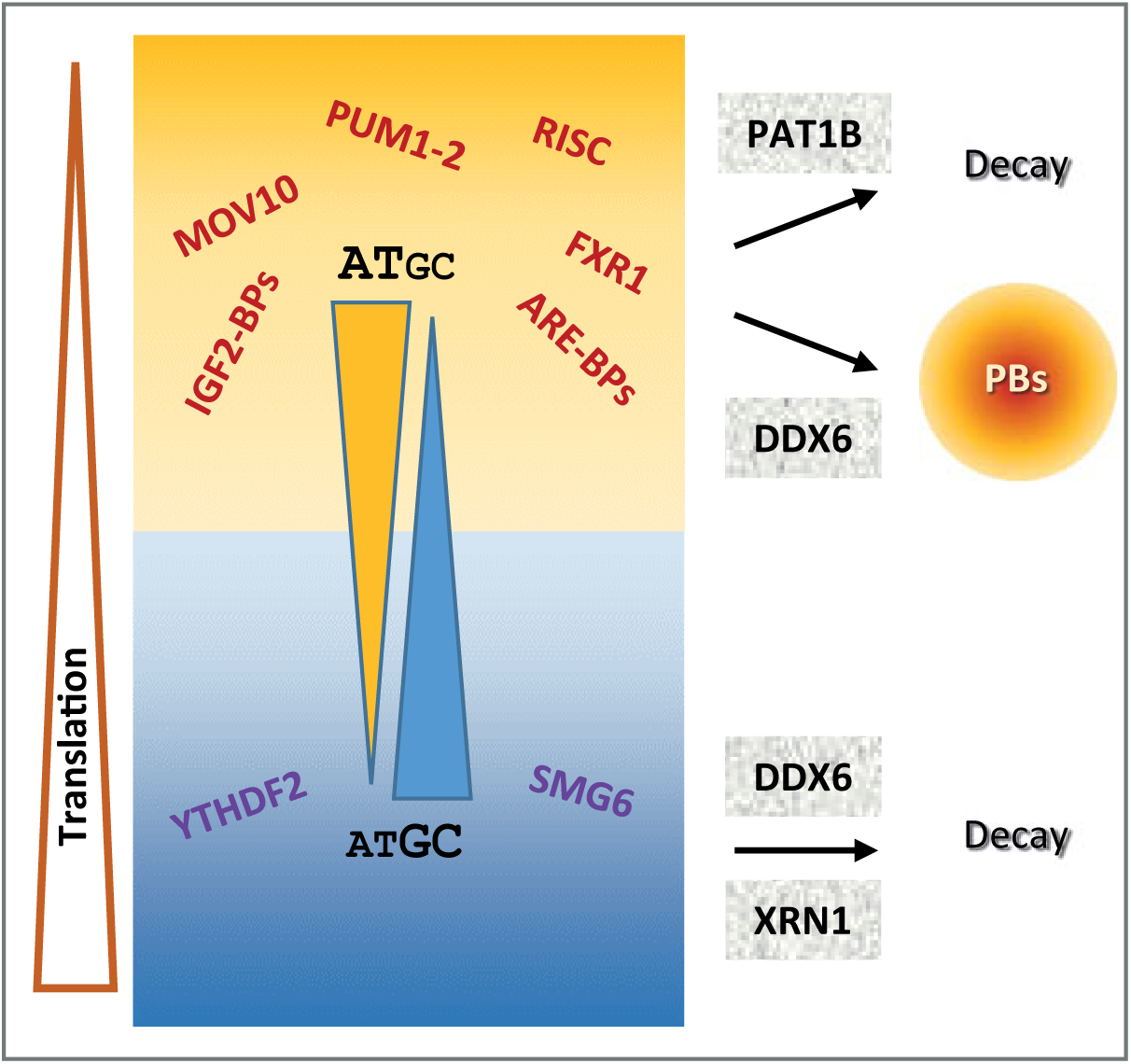
Schematic representation recapitulating the features of the mRNAs regulated at the level of decay and translation depending on their GC content.

It should be stressed that our analysis focused on trends common to most transcripts, which does not preclude that particular mRNAs could be exceptions to this general model, being GC-rich and translationally controlled, or AU-rich and regulated by 5’ decay. Moreover, while the analysis was consistent in proliferating cells of various origins, giving rise to a general model, it is possible that changes in cell physiology at particular developmental stages or during differentiation, rely on a different mechanism.

### GC content and codon usage

While the redundancy of the genetic code should enable amino acids to be encoded by synonymous codons of different base composition, the wide GC content variation between PB-enriched and PB-excluded mRNAs has consequences on the amino acid composition of encoded proteins. It also strongly impacts the identity of the wobble base: in PB mRNAs, the increased frequency of A/U at position 3 of the codon mechanically results in an increased use of low usage codons. As CDS are also longer in PB mRNAs, it further increases the number of low usage codons per CDS in these mRNAs. Interestingly, we showed that the absolute number of low usage codons per CDS best correlates with low protein yield. Thus, these results provide a molecular mechanism to a previously unexplained feature of PB mRNAs, that is, their particularly low protein yield, which we reported was an intrinsic property of these mRNAs and not simply the result of their sequestration in PBs [2]. Interestingly, the mRNAs of haplo-insufficiency genes, which by definition are expected to have a limited protein yield, are indeed enriched in PBs (Figure 6D).

In addition to the GC-dependent codon bias, we also observed some GC-independent codon bias in PB-enriched mRNAs. Interestingly, some important post-transcriptional regulation programs involve codon usage. This was shown for proliferation-versus differentiation-specific transcripts in human cells [46] and for maternal mRNA clearance during the maternal-to-zygotic transition in zebrafish, *Xenopus*, mouse, and *Drosophila* [47]. Codon usage could also enable the regulation of small subsets of mRNAs, depending on cellular requirements. In man, half of GO sets show more variability in codon usage than expected by chance [48]. Based on GO analysis, we previously demonstrated that PB mRNAs tend to encode regulatory proteins, while PB-excluded mRNAs encode basic functions. Furthermore, proteins of the same complexes tended to be encoded by mRNA co-regulated in or out of PBs, in so-called post-transcriptional regulons [2,3]. We speculate that specific codon usage could also underlie these post-transcriptional regulons.

### Distinct decay pathways

Our analysis distinguished separate decay pathways, depending on mRNA GC content. Interestingly, our previous analysis of the read coverage of PB and non-PB RNAs also suggested the existence of distinct decay pathways related to 3’ and 5’ extremities, respectively [2]. Part of the triage towards the PAT1B versus the DDX6/XRN1 pathway could be somehow associated with the capacity of the transcripts to condense in PBs, since, among the AU-rich mRNAs, only the ones enriched in PBs were susceptible to PAT1B-dependent decay (not shown). Nevertheless, PBs do not directly mediate the triage, since (i) TOP mRNAs were strongly excluded from PBs, but unaffected by either DDX6 or XRN1 silencing, and (ii) while PBs disappeared after DDX6 silencing [5], causing the release of AU-rich mRNAs into the cytosol, only GC-rich mRNAs were stabilized.

Focusing on DDX6-mediated decay, the minor impact of the 5’UTR GC content compared to CDS and 3’UTR, indicates that this helicase is not simply involved in allowing XRN1 access to the 5’ end. It is tempting to propose rather that, by unwinding GC-rich double-stranded regions over the entire length of the mRNA, DDX6 facilitates XRN1 progression. UPF1, another RNA helicase involved in mRNA decay, has also been shown to preferentially affect the decay of GC-rich mRNAs [49]. The same observation was made for targets of the NMD pathway, which involves UPF1, SMG6 and SMG7 [50]. Although the bias in these cases was restricted to the 3’UTR regions, it suggests that DDX6 could act in concert with other helicases for decay of GC-rich mRNAs. For AU-rich mRNAs, either such an active unfolding would be dispensable, or it would rely on other helicases that remain to be identified, with potential candidates being those enriched in purified PBs [2].

Concerning XRN1, we showed that it preferentially acts on PB-excluded mRNAs, that is, on GC-rich mRNAs but also, to a lesser extent, on a subset of AU-rich mRNAs (Figure S12A). While XRN1 is described as a general decay factor, there is evidence that 5’ decay shows some specificity *in vivo*. Indeed, in *Drosophila*, mutations in XRN1 have specific phenotypes, including wound healing, epithelial closure and stem cell renewal in testes, suggesting that it specifically degrades a subset of mRNAs [51]. Most importantly, a recent study showed that yeast XRN1 associates with ribosomes and decays mRNAs during translation [52]. If the mechanism is conserved in human, it would explain why XRN1 preferentially acts on GC-rich mRNAs, since they are the most actively translated mRNAs.

Turning to PAT1B, we showed that its silencing did not affect the same mRNAs as DDX6 silencing. A recent yeast study also showed that Dhh1/DDX6 and Pat1/PAT1B decay targets poorly overlap, suggesting the existence of two separate pathways in yeast as well [38]. However, the underlying mechanisms may differ in the two organisms, since both PAT1B and DDX6 decay targets were poorly translated in yeast, while only PAT1B targets were poorly translated in our study. Previous studies reported that tethered PAT1B decreases the abundance of a reporter mRNA in human cells, as a result of enhanced deadenylation and decapping [27,53,54]. Here, we did not find any evidence of PAT1B-dependent 5’ to 3’ decay in our read coverage analysis. Interestingly, genome-wide evidence in yeast too suggests that following decapping a significant fraction of the transcripts up-regulated in cells lacking Pat1 or Lsm1 is efficiently decayed 3’-5’, rather than by the 5’-3’ Xrn1 exonuclease [38]. Moreover, CLIP experiments in yeast showed a preference for Pat1 and Lsm1 binding to the 3’ end of mRNAs [55]. Altogether, we therefore favor the possibility that the mechanism of PAT1B-dependent decay is prominently 3’ to 5’ in human cells. It could involve the CCR4/CNOT deadenylase and the LMS1-7 complexes, as, despite their low abundance or small size, CNOT1 and LSM2/4 had high scores in our previous PAT1B interactome analysis [9,28].

PAT1B showed a strong preference for AU-rich targets, including those containing AREs. Many studies have demonstrated a link between AREs and mRNA stability, and its striking importance for processes such as inflammation [20]. Most ARE-BPs promote mRNA destabilization while some ARE-BPs, such as HuR [56,57] and AUF1 for a subset of mRNAs [58], can stabilize mRNAs. Altogether, these observations raise the possibility that ARE-BPs behave either as enhancers or inhibitors of PAT1B activity in mRNA decay. Similarly, the miRNA pathway could activate PAT1B activity in decay.

### Translation repression and PB accumulation

DDX6 activity in translation repression has been documented in a variety of biological contexts. In *Xenopus* oocytes, DDX6 contributes to the repression of maternal mRNAs, as a component of the well characterized CPEB complex [18]. In *Drosophila*, Me31B/DDX6 represses the translation of thousands of mRNAs during the early stages of the maternal to zygotic transition [59]. It also collaborates with FMRP and AGO proteins for translation repression in fly neurons [60]. In mammals, DDX6 is a general co-factor of the miRNA pathway [13,14,16,41]. The intriguing finding of our analysis was that the targets of most tested translation regulators (FRX1-2, FMR1, PUM1-2, most miRNAs…) were AU-rich and had a median behavior in all datasets similar to other mRNAs of same GC content. A provocative possibility would be that these RBPs bind a large number of transcripts, but only affect a minority of them, depending on other associated RBPs. Alternatively, the specific activity of these RBPs could become significant only when their expression is turned on or off. Of note, while TOP mRNAs clearly constituted an exception in terms of GC content, they are particular too in terms of translation repression mechanism, with a regulating sequence located at the 5’ end.

PBs add another layer to translation regulation, by storing translationally repressed mRNAs. It was already known that ARE-containing mRNAs bound to ARE-BPs such as TTP and BRF were recruited to PBs [34] and that miRNA targets accumulate in PBs upon miRNA binding in a reversible manner [40,61]. As DDX6 and 4E-T are key factors in PB assembly in mammalian cells [5,6,53,62], it raises the question of whether these proteins contribute to translation repression by triggering the recruitment of arrested mRNA to PBs, or by mediating translation arrest, which then results in mRNA recruitment to PBs. While we have no answer for DDX6, we observed that 4E-T targets were particularly enriched in PBs, though rather insensitive to DDX6 or PAT1B knock-down. First, this suggests that 4E-T function in PBs is partly independent of DDX6, agreeing with the previous observation that some PBs can still form when the DDX6 interaction domain of 4E-T is mutated [16]. Second, it indicates that PB localization and translation repression by DDX6 can be separated, at least to some extent.

In addition to their high AU content, we observed that PB mRNAs were longer than mRNAs excluded from PBs. Long CDS could favor mRNA recruitment in PBs by decreasing translation efficiency, and hence increasing the fraction of polysome-free mRNAs. Long 3’UTR should increase the probability of binding translation regulators, contributing also to PB recruitment. In agreement, it is interesting that DDX6 preferentially repressed the translation of mRNAs with long 3’UTR, while the CDS length was irrelevant (Figure S4G). It is also possible that protein binding over the entire length of mRNA may contribute to PB recruitment. This would explain why the GC content of the 5’UTR has little impact compared to CDS and 3’UTR, as leader sequences, being considerably shorter, would bind less of such proteins. In this regard, it is interesting to note that we and others have previously proposed from biochemical experiments and electron microscopy imaging that DDX6 and its partner LSM14A coat repressed mRNAs at multiple positions, according to their length [25,63].

### Evolutionary issues

Our results raise intriguing issues in terms of evolution. While PBs are found in all eukaryotes, animal and vegetal, the GC-rich part of the human genome only emerged in amniotes (the ancestor of birds and mammals) [64]. In more distant organisms, such as yeast, *C. elegans*, *Drosophila* or *Xenopus*, genes have a narrow GC content distribution, most often AU-rich (Figure S13). Thus, despite the conservation of the DDX6, XRN1 and to a lesser extent PAT1B proteins in eukaryotes, distinct mRNA decay pathways depending on GC content may have evolved more recently. Moreover, the enzymatic properties of DDX6 could have adapted to the higher GC content of human transcripts.

The GC-rich part of the human genome was acquired through GC-biased gene conversion (gBGC), a non-selective process linked with meiotic recombination affecting GC content evolution [22]. We considered the possibility that meiotic recombination occurred more frequently in genomic regions containing genes involved in basic functions, leading to stronger gBGC and, consequently, to higher GC content of PB-excluded mRNA. However, our analysis showed that mRNA base composition and PB enrichment are associated independently of meiotic recombination or the genomic context. We therefore put forward a model where the genome of higher eukaryotes has evolved partly to facilitate the control of regulators at the translation level, by limiting their protein yield. Regardless, the overall outcome of our study is that in human the GC content, a feature written in the genome, shapes mRNA fate and its control in a strikingly coherent system.

## Materials and methods

### Cell culture and transfection

Human embryonic kidney HEK 293 cells and epithelioid carcinoma HeLa cells were maintained in DMEM supplemented with 10% (v/v) fetal calf serum. HCT116 were grown in McCoy’s 5A modified medium supplemented with 10% (v/v) fetal bovine serum, 5% (v/v) sodium pyruvate and 5% (v/v) non-essential amino acids.

For DDX6 silencing, 7.105 cells were transfected at the time of their plating (reverse transfection) with 50 pmoles DDX6 or control β-globin siRNAs [5] per 3 cm diameter well, using Lipofectamine 2000 (Life Technologies, France), and split in two 24 hours later. Cells were lyzed 48 h after transfection.

For XRN1 silencing with siRNAs, 2.105 cells/well were plated in 6-well plates and transfected 24 h later with 50 nM siRNA negative Control or Silencer Pre-designed siRNA XRN1, using Lipofectamine RNAiMAX (Life Technologies). Cells were lyzed 48 h after transfection.

For XRN1 silencing with shRNA, a doxycycline inducible construct provided by Dharmacon (TRIPZ) with shRNA against XRN1 or non-silencing shRNA was introduced by lentiviral transduction (MOI 0.5). After 10 days of puromycin selection, cells were tested for expression of the construct. For shRNA induction cells were grown to 30% confluency in 10 cm plates before adding 1 µg/ml doxycycline. After 24h, cells were split in three and doxycycline was maintained until 48h.

### Western blot analysis

Total cell lysates were obtained as described previously [65]. Proteins were separated on SDS-PAGE on 4–12% polyacrylamide gel (NuPage, Invitrogen) and transferred onto nitrocellulose membrane (PerkinElmer, France). After blocking in 5% (w/v) nonfat dry milk in PBS for 30 min at room temperature, the membrane was incubated for 1 h at 37°C with primary antibodies. After washing in PBS containing 0.05% (v/v) tween-20, blots were incubated for 40 min at room temperature with horseradish peroxidase-conjugated secondary anti-rabbit antibody (1:10000; Jackson Immunoresearch Laboratories). Immunoreactive bands were visualized by chemiluminescence detection of peroxidase activity (SuperSignal West Pico, Pierce) and exposure to CL-XPosure film (Pierce). Protein expression was evaluated by densitometry (NIH ImageJ).

For the shRNA XRN1 experiment, proteins were isolated using RIPA buffer with Halt™ Protease and Phosphatase Inhibitor Cocktail (Thermo Scientific), precipitated with acetone and separated on Tris-Acetate 3-8% polyacrylamide gel (NuPage, Invitrogen) before transfer to nitrocellulose membrane (GE Healthcare). After blocking with 5% (w/v) nonfat dry milk in TBST for 30 min at room temperature the membrane was incubated for 1h at room temperature or o/n at 4°C with primary antibodies. After washing in TBST, blots were subsequently incubated for 1h at room temperature with horseradish peroxidase-conjugated secondary anti-rabbit/mouse antibodies (1:10000; Sigma). Immunoreactive bands were visualized by chemiluminescence detection of peroxidase activity (SuperSignal West Dura, Pierce) and imaged using ImageQuant LAS 4000 (GE Healthcare). Protein expression was evaluated by densitometry (NIH ImageJ).

Primary antibodies were: rabbit polyclonal anti-DDX6 (1:15000; Novus Biological), rabbit polyclonal anti-ribosomal S6 (1:5000; Cell Signaling Technology), anti-XRN1 (1:1000, Bethyl Laboratories), anti-XRN1 (1:5000 Novus Bioscience), anti-Pol II (1:100, Santa Cruz).

### Polysome profiling experiment

The polysome profiling experiment was described in [2]. For q-RT-PCR experiments, 1 μg RNA was reverse transcribed for 1 h at 50°C using the SuperScript III First-Strand Synthesis System for RT-PCR (Life Technologies) with 1 μg random primers. No amplification was detected in negative controls omitting the reverse transcriptase. qPCR amplifications (12 μl) were done in duplicates using 1 μl of cDNA and the GoTaq Probe 2X Master Mix (Promega) on a LightCycler 480 (Roche), programmed as follows: 5 min, 95°C; 40 cycles [10 sec, 95°C; 15 sec, 60°C; 10 sec, 72°C]. A last step (5 min progressive 95°C to 72°C) calculating the qPCR product Tm allowed for reaction specificity check. Primers for ACTB, APP, BACE1, LSM14A, LSM14B, MFN2, PNRC1, TIMP2 and TRIB1 were designed using the Primer 3 software [66]. The results were normalized using HPRT1.

### Library preparation and RNA-Seq data processing

For polysome profiling after DDX6 silencing and transcriptome after PAT1B silencing in HEK293 cells, rRNA was depleted using the Ribo-Zero kit Human/Mouse/Rat (Epicentre), and libraries were prepared using random priming. Triplicate and duplicate libraries were generated from three or two independent experiments, respectively, and processed as detailed previously [2,9].

For the transcriptome after XRN1 silencing in HeLa cells, libraries were prepared from 500 ng of total RNAs and oligo(dT) primed using TruSeq Stranded Total RNA kit (Illumina) with two technical replicates for each sample. Libraries were then quantified with KAPA Library Quantification kit (Kapa Biosystems) and pooled. 4 nM of this pool were loaded on a high output flowcell and sequenced on a NextSeq500 platform (Illumina) with 2×75nt paired-end chemistry.

For the shRNA XRN1 experiment RNA was isolated using Quiazol and miRNase Mini Kit (Quiagen), next subjected to DNase treatment (Quiagen) and quality control with Bioanalyzer. The rRNA was removed using rRNA Removal Mix. Libraries were prepared from 1 ug of RNA following TruSeq Stranded Total RNA kit (Illumina) with two technical replicates for each sample. 100nt paired-end RNA-Seq was performed on HiSeq - Rapid Run (Illumina). The results were aligned using hg19 genome and DESeq2, with standard settings, was used for determining FC and p-values.

For PB enrichment, libraries were prepared without prior elimination of rRNA and using random priming. Triplicate libraries were generated from three independent experiments and processed using the same pipeline as for DDX6 silencing [2].

For the transcriptome after induction of a stably transfected DDX6 shRNA for 48h in K562 cells, the .fastq files from experiments ENCSR119QWQ (DDX6 shRNA) and ENCSR913CAE (control shRNA) were processed according to the same pipeline as DDX6 polysome profiling, except that the control and DDX6 shRNA experiments were not paired to compute the corrected p-values of RNA differential expression in EdgeR_3.6.2.

The ENCODE dataset of mRNA clipped to DDX6 in K562 cells was generated using ENCODE .bam files aligned on the hg19 genome corresponding to (i) the DDX6 eClip experiment ENCSR893EFU, and (ii) the total RNA-seq of K562 experiment ENCSR109IQO. The enrichment in the CLIP dataset compared to the total RNA sample was calculated as in the DDX6 shRNA experiment.

### Bioinformatic analysis

Briefly, the read coverage was computed as follows. Raw reads were processed using trimmomatics. Alignment was performed on the longest transcript isoforms of Ensembl annotated genes with bowtie2 aligner. Isoforms shorter than 500 nucleotides were not considered. Only unique mapped reads were qualified for counting. Each transcript was subdivided in 10 bins from transcription start site (TSS) to transcript end site (TES) and the proportion of reads for each bin was computed. For the metagene analysis, the average distribution of read along transcript length was computed so that each gene had the same weight independently of it expression level.

The protein yield was calculated as the ratio between protein abundance in HEK293 cells, taken from [67], and mRNA abundance in HEK293 cells, taken from the control sample of our DDX6 polysome profiling experiment.

For GC profiling of the transcripts in various organisms (Figures 1B and S13), transcripts were downloaded from ENSEMBL (version 92) with their associated GC content.

Boxplot representations and statistical tests were performed using the GraphPad Prism software (GraphPad software, Inc.) and the R suite (https://www.R-project.org) (R Core Team 2018. R: A language and environment for statistical computing. R Foundation for Statistical Computing, Vienna, Austria). Other graphical representations were generated using Excel and the Excel Analysis ToolPak (Microsoft). Hierarchical clustering of all transcripts in Figure 4E was performed using the Cluster 3.0/Treeview softwares (Kendall’s tau distance, average linkage, [68]. Heatmap representation of the targets of the various regulators in Figures 5B and D was performed online using Morpheus (https://software.broadinstitute.org/morpheus).

Gene meiotic recombination rates were computed as crossover rates between gene start and gene end using the genetic map from the HapMap project [69]. Rates were computed as the weighted average of crossover rates of chromosomal regions that overlap the window.

### Datasets used in the bioinformatics analysis

The following datasets were downloaded from the supplementary material of the corresponding papers: 1) For mRNA-containing cis-regulatory motifs: (i) in silico identification: AREs [70]; (ii) experimental determination: CPEs [71], G4-containing [72] and TOP mRNAs [33]. G4-containing genes were restricted to those harboring a G4 in 5’UTR. 2) For RBP targets: (i) PARE in HeLa cells: SMG6 (mRNAs actually cleaved by SMG6) [29] (ii) CLIP in HEK293 cells: HuR and TTP [57,73]; (iii) CLIP in HeLa cells: YTHDF2 [30]; (iv) RIP-CHIP in HeLa cells: PUM1 [74]; (v) RIP-CHIP in neurons: 4E-T [75]. In the case of HuR, the transcripts clipped only in 5’UTR/CDS/introns were removed, and the target list was restricted to transcripts clipped more than once. In the case of TTP, the transcripts clipped only in introns were removed.

For other RBP targets (ATXN2, MOV10, IGF2BP1-3, PUM2, FMR1, FXR1-2, AGO1-4, YTHDF2), CLIP data from different laboratories were previously processed through the same pipeline in the CLIPdb 1.0 database using the Piranha method [31]. We retained those performed in epithelial cells (HeLa, HEK293, HEK293T). Moreover, when replicates were available, we selected the RNA-protein interactions detected in at least 50% of the replicates. Except for FXR1-2 targets, we determined whether protein-RNA interactions occurred in UTR or CDS by intersecting coordinates of the read peaks with the v19 gencode annotation, and we removed the transcripts clipped in their CDS.

For miRNA targets, we extracted the list of all experimentally documented targets from miRTarBase (http://mirtarbase.mbc.nctu.edu.tw/php/index.php) [43], and selected the targets of the 22 miRNAs of interest.

### Data availability

RNA-Seq gene expression data and raw fastq files are available on the SRA repository https://www.ebi.ac.uk/ena under accession number: E-MTAB-4091 for the polysome profiling after DDX6 silencing, E-MTAB-5577 for the transcriptome after PAT1B silencing, E-MTAB-5477 for the PB transcriptome.

RNA-Seq gene expression data from XRN1 silencing experiments is available on the GEO repository: GSE115471 and GSE114605 for the HeLa and HCT116 datasets, respectively.

All ENCODE experiments are available at https://www.encodeproject.org.

## Declarations

### Competing interests

The authors declare that they have no competing interests.

### Authors’ contributions

All authors read and approved the final manuscript.

## Acknowledgments

We thank Michèle Ernoult-Lange, Marina Pinskaya and Marc Gabriel for scientific discussions and technical assistance. We also thank Virginie Magnone, Kevin Lebrigand (NGS platform, UCA Genomix), Sylvain Baulande, Patricia Legoix-Né and Virginie Raynal (NGS platform, Institut Curie).

## Funding

This work was supported by the Association pour la Recherche sur le Cancer and the Agence Nationale pour la Recherche contract ANR-14-CE09-0013-01 to D.W.’s laboratory, the European Research Council (DARK consolidator grant) to A.M.’s laboratory, the LABEX SIGNALIFE [ANR-11-LABX-0028-01], Canceropole PACA to P.B.’s laboratory, BBSRC, Newton Trust and Foundation Wiener – Anspach (C.V.) to N.S.’s laboratory. This work has benefited from the facilities and expertise of the NGS platform of Institut Curie (supported by the Agence Nationale de la Recherche [ANR-10-EQPX-03, ANR10-INBS-09-08] and the Canceropôle Ile-de-France), and of the “UCA GenomiX” Functional Genomics Platform of the University of Nice Sophia Antipolis, a partner of the National Infrastructure France Génomique, supported by Commissariat aux Grands Investissements (ANR-10-INBS-09-03, ANR-10-INBS-09-02) and Canceropôle PACA.

## Notes

#### Summary of Updates

The manuscript has been rewritten for easier understanding. Moreover, several analysis have been added to clarify the respective contribution (i) of RNA GC content and length, and (ii) of CDS and 3'UTR.

