## Supplemental figures for "GC content shapes mRNA decay and storage in human cells"

#### Supplementary information

**Figure S1: The GC content is more important for PB than SG localization.**
(A) Number of transcripts in the different classes analyzed in Figures 1A,E.
(B) Number of transcripts in the different classes analyzed in Figures 1B,C and S1B,C.
(C) Stress granule localization is weakly dependent on GC content. The SG transcriptome analysis from [21] was analyzed as in Figure 1A.
(D) PB-enriched mRNAs are longer than PB-excluded ones independently of their GC content. The distribution of length of PB-enriched (PB-in) and PB-excluded (PB-out) mRNAs was analyzed as in Figure 1E.

**Figure S2: Codon usage biases.**
(A) PB mRNAs and PB-excluded mRNAs encode proteins with different amino acid composition. The boxplots represent the distribution of the frequency of each amino acid in the proteins encoded by mRNAs enriched (PB-in, in red, n=5500) or excluded from PBs (PB-out, in blue, n=5104). The boxes represent the 25-75 percentiles and the whiskers the 10-90 percentiles.
(B,C) Codon usage biases independent on the GC content. For each amino acid with 4 or 6 synonymous codons, the ratio of usage of NNU to NNA (A) or NNC to NNG (B) was calculated. The boxplots represent the distribution of these ratios in PB-enriched (PB-in) and PB-excluded (PB-out) mRNAs using a PB enrichment threshold of +/- 1 in log2 scale. All differences were statistically significant (*, p-value <0.05) except for the Serine NNT/NNA ratio. Note the change of scale between (A) and (B).
(D) Low usage codons are more frequent in PB mRNAs. The ratio between the relative codon usage in PB-enriched and PB-excluded (PB-in/PB-out) mRNAs was expressed as a function of their global relative usage, after normalization by the number of synonymous codons.

**Figure S3: Polysome profiling following DDX6 silencing.**
(A) Western blot analysis of DDX6 in siRNA-transfected HEK293 cells. The DDX6 signal, normalized using the ribosomal protein S6, is indicated on the right panel (n=3; p<0.001; t-test).
(B) Representative polysome profiles of siRNA-transfected cells.
(C,D) MA plots of mRNA fold changes between siRNA-transfected and control cells in total (C) and polysomal (D) mRNA.
(E) Correlation between fold changes (FC) of randomly selected mRNAs measured by qPCR and calculated from RNA-Seq data, in total and polysomal fractions.
(F,G) Correlation between total (F) and polysomal (G) mRNA fold-changes and the GC content of their gene. Note that correlation is positive for total mRNAs and negative for polysomal mRNAs. rs is the Spearman correlation coefficient.

**Figure S4: Impact of DDX6 binding and mRNA length on DDX6 dependency.**
(A) Number of transcripts in the different classes analyzed in Figures 3 and 4A,B.
(B) Dot plot representation of the changes in translation rate (Polysome/Total FC) as a function of changes in mRNA stability (Total FC) after DDX6 silencing.
(C,D) DDX6-clipped targets tend to be particularly stabilized after DDX6 silencing in both HEK293 and K562 cells (C), but less affected than other mRNAs in terms of translation rate (D).
(E) Number of transcripts in the different classes analyzed in (F) and (G).
(F) Transcript length has a weak impact on mRNA stabilization following DDX6 silencing. Transcripts were subdivided into six classes depending on their full length (from <1.5 to 10 kb, left panel), the length of their CDS (from <0.5 to 6 kb, middle panel) or their 3'UTR (from <0.5 to 6 kb, right panel). The boxplots represent the distribution of their respective fold-changes in total mRNA. The difference

between class 1 and 6 was statistically significant using a two tail Mann-Whitney test: p<0.0001 in all panels. rs is the Spearman correlation coefficient.
(G) The 3'UTR length has an impact on mRNA translation derepression following DDX6 silencing. The data are represented as in (F). Note that the difference between class 1 and 6 was statistically significant in left and right panels only.

**Figure S5: Impact of the GC content on DDX6-dependency.**
(A) Number of transcripts in the different classes analyzed in (B) and (C).
(B,C) mRNA stabilization (B) and translation derepression (C) following DDX6 silencing in HEK293 cells mostly depends on the GC content of their CDS and 3' UTR. The analysis was performed as in Figure 1C.

**Figure S6: Transcriptome analysis following XRN1 and PAT1B silencing.**
(A) Western blot analysis of XRN1 in siRNA-transfected HeLa cells. The XRN1 signal, normalized using tubulin, is indicated below.
(B) Western blot analysis of XRN1 in shRNA-transfected HCT116 cells. Cells transfected with the XRN1 or control shRNA were induced for 0 to 72 h with Doxycycline. The XRN1 signal, normalized using RNA Pol II, is indicated below. The time point 48 h was chosen for library preparation.
(C) XRN1 targets tend to be excluded from PBs. The fold-changes after XRN1 silencing in HeLa cells were expressed as a function of PB enrichment.
(D, E) PAT1B targets are regulated by DDX6 at the level of translation but not decay. The total (D) and polysomal (E) mRNA fold-changes (FC) after DDX6 silencing were expressed as a function of fold-changes after PAT1B silencing.
(F) PAT1B targets are enriched in PBs. The fold-changes (FC) of total RNA after PAT1B silencing were expressed as a function of PB enrichment.
For each panel, rs is the Spearman correlation coefficient.

**Figure S7: Targets of group I and II regulators (part I).**
(A) Number of transcripts in the mRNA subsets analyzed in (B-D) and in Figures 5A,B, S8A-D and S10.
(B) Sensitivity of the different mRNA subsets to DDX6 silencing in HEK293 (upper panel) and K562 cells (lower panel) in terms of decay. The behavior of all mRNAs is shown for comparison (in grey). Only the targets of group I factors were stabilized. While the presented analysis uses the YTHDF2 targets reported in [31], the same pattern was observed using the targets reported in [30].
(C) Sensitivity of the different mRNA subsets to XRN1 silencing in HeLa (upper panel) and HCT116 cells (lower panel). Only the targets of group I factors were stabilized in HCT116 cells, while only SMG6 targets were stabilized in HeLa cells.
(D) Sensitivity of the different mRNA subsets to PAT1B silencing in HEK293 cells. Only the targets of group II factors were stabilized, except ATXN2.
Two-tail Mann-Whitney test was performed with respect to all mRNAs. **, p<0.0001; ns, non-significant.

**Figure S8: Targets of group I and II regulators (part II).**
(A) PB localization of the different mRNA subsets. Only the targets of group II factors were enriched in PBs, except ATXN2. Note also that SMG6 targets and TOP mRNAs were particularly excluded from PBs.
(B) Sensitivity of the different mRNA subsets to DDX6 silencing in HEK293 in terms of translation. Only the targets of group II factors were translationally derepressed, except ATXN2.
Two-tail Mann-Whitney test was performed with respect to all mRNAs. **, p<0.0001.

(C, D) The targets of group I regulators were stabilized after DDX6 (C) and XRN1 (D) silencing mostly like other PB-excluded mRNAs. The same analysis as in Figure S7B and S7C was conducted separately on PB-excluded (PB-out) and PB-enriched (PB-in) mRNAs. The distribution of all mRNAs (in grey), all PB-out and all PB-in mRNAs (dashed boxes) are shown for comparison. Two-tail Mann-Whitney test was performed with respect to cognate PB-out or PB-in mRNAs. **, $p<0.0001$; *, $p=0.006$ for (C) and 0.02 for (D); ns, non-significant.
(E) DDX6-dependent mRNA decay does not correlate with polysome engagement. The fraction of mRNAs in polysomes (polysomal/total cpm) was calculated using the RNA-Seq data from control HEK293 cells. Transcripts were subdivided into 15 classes (1000 transcripts each) of increasing value. The boxplot represents their fold-change (FC) distribution in total RNA following DDX6 silencing.

**Figure S9: Targets of the miRNA pathway.**
(A,B) Number of AGO targets (A) and miRNA targets (B) analyzed in (C-H) and in Figures 5C,D.
(C-E) AGO targets are enriched in PBs (C), translationally derepressed (D) and little or not stabilized (E) following DDX6 silencing. Two-tail Mann-Whitney test was performed with respect to all mRNAs. **, $p<0.0001$; *, $p=0.002$, x, $p=0.04$; ns, non-significant.
(F) AGO targets excluded from PBs are stabilized following DDX6 silencing like other PB-excluded transcripts. The analysis was performed as in Figure S8C. Two-tail Mann-Whitney test was performed with respect to cognate PB-out or PB-in mRNAs. x, $p=0.04$; ns, non-significant.
(G) AGO targets are stabilized following PAT1B silencing. Two-tail Mann-Whitney test was performed with respect to all mRNAs. **, $p<0.0001$; x, $p=0.04$.
(H) The extent of PB enrichment of miRNA targets depends on the miRNA. Two-tail Mann-Whitney test was performed with respect to all mRNAs. **, $p<0.0001$; ns, non-significant.

**Figure S10: Targets of the group I and II regulators behave like mRNAs of similar GC content.**
(A-E) The whole transcriptome was subdivided in groups of 500 mRNAs after ranking depending on the GC content of their gene. The median GC content of each group was represented as a function of their median fold-change in the various datasets (in grey). The same analysis was performed for the targets of group I (in green) and II (in orange) regulators, and superimposed to the graphs. The dashed line indicates the median GC content for all mRNAs. Overall, in all datasets, the targets of the different regulators behave similarly to mRNAs of same GC content. For figure clarity, only the few regulators leading to distinct behaviors were indicated.

**Figure S11: miRNA targets behave like mRNAs of similar GC content.**
(A-E) The analysis was conducted as in Figure S10. Overall, the targets of the different miRNAs behave similarly to mRNAs of same GC content.
(F) The GC content of miRNA targets is correlated with the GC content of the miRNA itself. rp is the Pearson correlation coefficient.

**Figure S12: Role of PB localization in XRN1 and DDX6 sensitivity and importance of the coding property for PB localization.**
(A) AU-rich mRNAs are better XRN1 targets when excluded than when enriched in PBs. The analysis was conducted as in Figure S10B for all mRNAs (in grey) and for PB-excluded mRNAs (PB-out, in dark blue), except that mRNAs with GC values under 50% were grouped by 200 to account for their low number.
(B) AU-rich mRNAs are not better DDX6 decay targets when they are excluded from PBs. The dataset after DDX6 silencing in HEK293 cells was analyzed as in (A).

(C) LncRNAs poorly accumulate in PBs even when they are AU-rich. LncRNAs (n=2589) were subdivided in 6 classes of increasing GC content and their accumulation in PBs was represented as for mRNAs in Figure 1B. rs is the Spearman correlation coefficient.

(D) Number of transcripts in the different classes analyzed in (C) and in Figures 6A,B.

**Figure S13: Distribution of the gene GC content in various eukaryotic genomes.**
The graphs were constructed as in Figure 1C. The median value for each genome is indicated below.

**Table S1: transcriptome datasets.**
Sheet1: polysome profiling after siDDX6 in HEK293 cells.
Sheet2: transcriptome after shDDX6 in K562 cells.
Sheet3: DDX6 CLIP in K562 cells.
Sheet4: transcriptome after siXRN1 in HeLa cells.
Sheet5: transcriptome after shXRN1 in HCT116 cells.
Sheet6: transcriptome after siPAT1B in HEK293 cells.

#### Figure S1

### A

| mRNA length (kb) | <1.5 | 1.5-2 | 2-3 | 3-5 | 5-10 | >10 |
|---|---|---|---|---|---|---|
| HEK293 / PBs (Fig 1A) | 2618 | 1866 | 3424 | 3822 | 2385 | 381 |
| PB-in (Fig 1E) | 407 | 424 | 1076 | 1793 | 1457 | 224 |
| PB-out (Fig 1E) | 1441 | 827 | 1300 | 979 | 399 | 68 |

### B

| GC (%) | <40 | 40-45 | 45-50 | 50-55 | 55-60 | >60 | 60-65 | 65-70 | >70 |
|---|---|---|---|---|---|---|---|---|---|
| HEK293 / PBs (Fig 1B) | 3967 | 4067 | 2739 | 2437 | 1595 | 843 | | | |
| PBs_5'UTR (Fig 1D) | 471 | 536 | 782 | 1326 | 1643 | | 2175 | 2276 | 4617 |
| PBs_CDS (Fig 1D) | 913 | 2669 | 2623 | 2372 | 2536 | 3377 | | | |
| PBs_3'UTR (Fig 1D) | 6010 | 2074 | 1758 | 1701 | 1455 | 1051 | | | |
| PB-in (Fig S1D) | 1485 | 1922 | 1168 | 572 | 189 | 44 | | | |
| PB-out (Fig S1D) | 114 | 349 | 573 | 934 | 1374 | 1670 | | | |
| SG enrichment (Fig S1C) | 2824 | 2972 | 1910 | 1710 | 1159 | 633 | | | |

### C

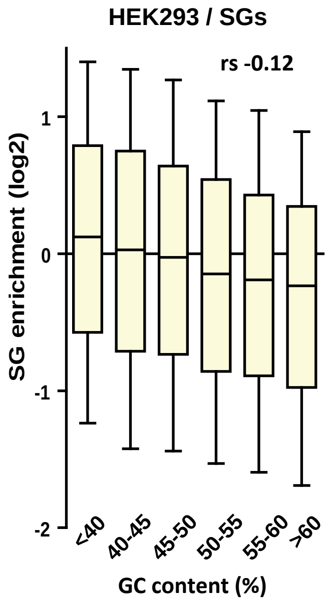


### D

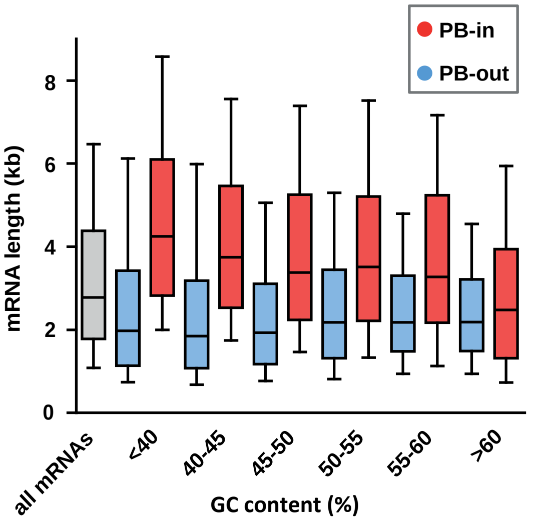

### Figure S2

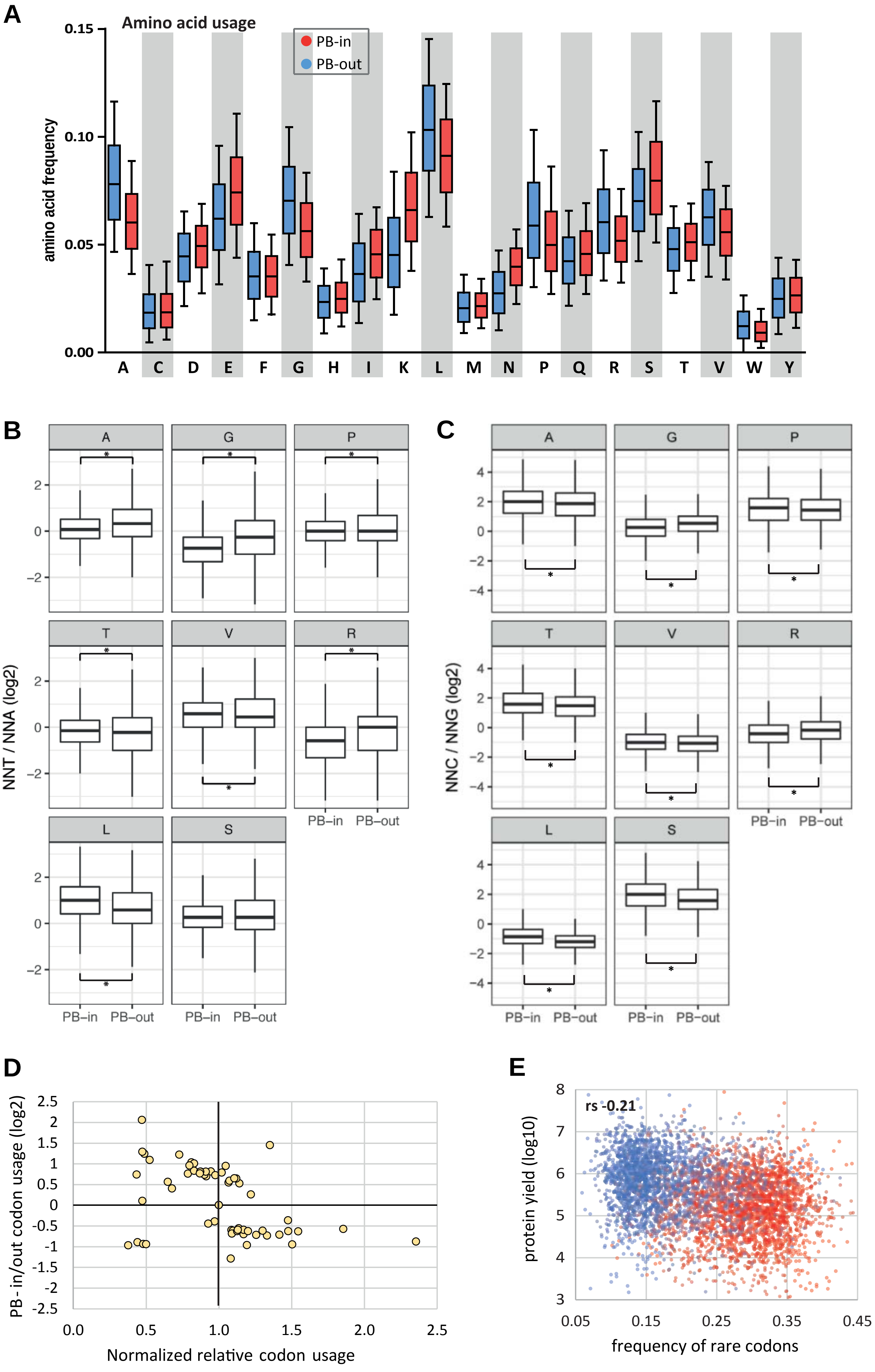
A
Amino acid usage
PB-in
PB-out
amino acid frequency
0.15
0.10
0.05
0.00
A C D E F G H I K L M N P Q R S T V W Y
B
NNT / NNA (log2)
A G P T V R L S
PB–in PB–out
C
NNC / NNG (log2)
A G P T V R L S
PB–in PB–out
D
PB - in/out codon usage (log2)
Normalized relative codon usage
E
rs -0.21
protein yield (log10)
frequency of rare codons

#### Figure S3

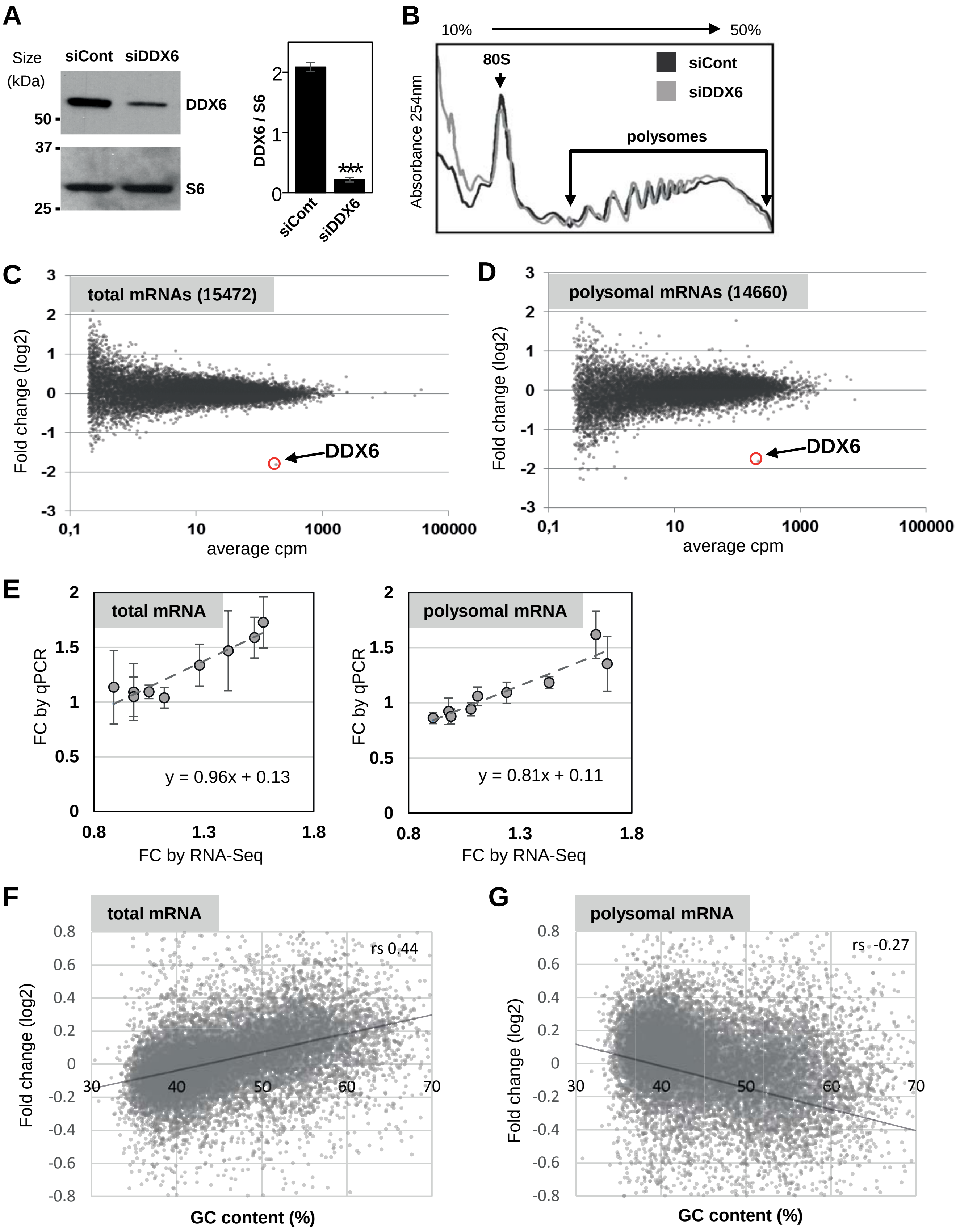
A
Size (kDa)
siCont siDDX6
50
37
25
DDX6
S6
DDX6 / S6
2
1
0
***
siCont
siDDX6
B
10%
50%
80S
siCont
siDDX6
polysomes
Absorbance 254nm
C
total mRNAs (15472)
Fold change (log2)
DDX6
average cpm
0,1
10
1000
100000
D
polysomal mRNAs (14660)
Fold change (log2)
DDX6
average cpm
E
total mRNA
FC by qPCR
y = 0.96x + 0.13
FC by RNA-Seq
polysomal mRNA
y = 0.81x + 0.11
FC by RNA-Seq
F
total mRNA
rs 0.44
Fold change (log2)
GC content (%)
G
polysomal mRNA
rs -0.27
Fold change (log2)
GC content (%)

### Figure S4

**A**

| GC (%) | <40 | 40-45 | 45-50 | 50-55 | 55-60 | >60 |
|---|---|---|---|---|---|---|
| HEK293 / siDDX6 (Fig 3A,C) | 3497 | 3590 | 2275 | 2065 | 1362 | 725 |
| K562 / shDDX6 (Fig 3B) | 2710 | 2835 | 1838 | 1778 | 1210 | 626 |
| K562 / CLIP DDX6 (Fig 3D) | 2901 | 3015 | 1973 | 1846 | 1228 | 712 |
| HeLa / siXrn1 (Fig 4A) | 2908 | 2948 | 1818 | 1669 | 1101 | 594 |
| HCT116 / shXrn1 (Fig 4A) | 3318 | 3507 | 2380 | 2251 | 1520 | 890 |
| HEK293 / siPat1b (Fig 4B) | 2901 | 3015 | 1973 | 1846 | 1229 | 712 |

**B**

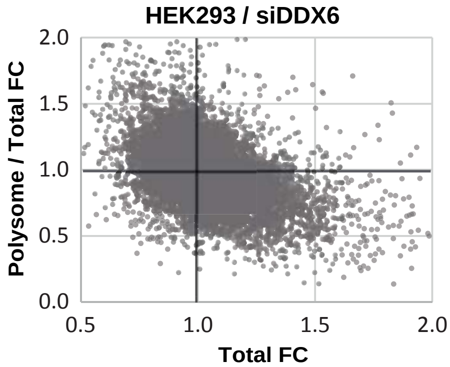


**C**

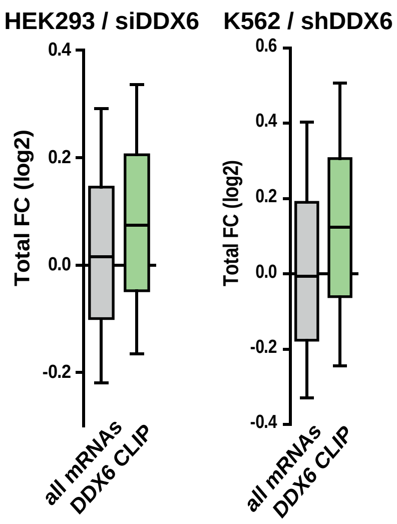


**D**

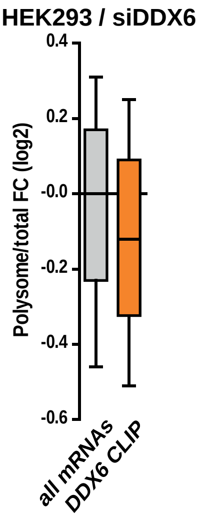


**E**

| mRNA length (kb) | | | | | |
|---|---|---|---|---|---|
| <1.5 | 1.5-2 | 2-3 | 3-5 | 5-10 | >10 |
| 1857 | 1583 | 3073 | 3779 | 2523 | 459 |
| **CDS length (kb)** | | | | | |
| <0.5 | 0.5-1 | 1-2 | 2-4 | 4-6 | >6 |
| 882 | 2929 | 5220 | 3201 | 632 | 401 |
| **3'UTR length (kb)** | | | | | |
| <0.5 | 0.5-1 | 1-2 | 2-4 | 4-6 | >6 |
| 4390 | 2448 | 2842 | 2350 | 771 | 469 |

**F**

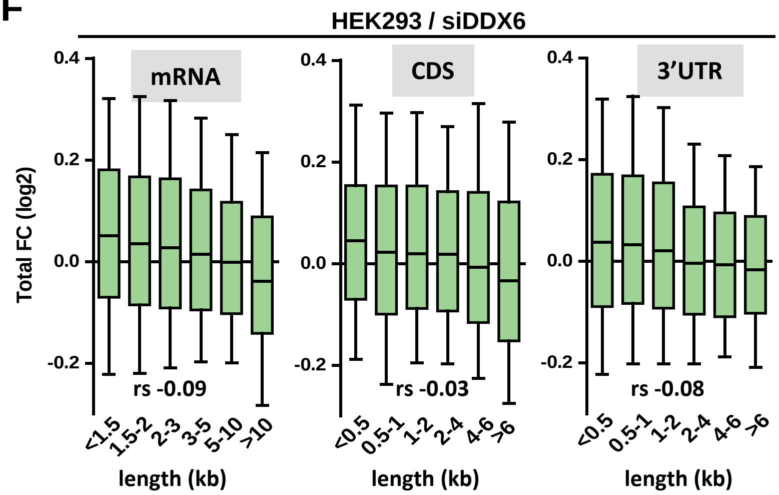


**G**

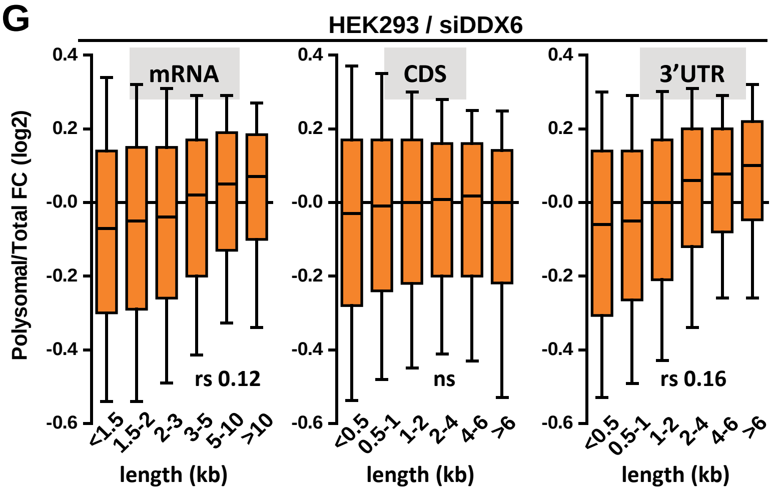

### Figure S5

**A**

| | | | | | |
|---|---|---|---|---|---|
| 5'UTR GC (%) | | | | | |
| <40 | 40-45 | 45-50 | 50-55 | 55-60 | >60 |
| 297 | 446 | 734 | 1207 | 1646 | 8371 |
| CDS GC (%) | | | | | |
| <40 | 40-45 | 45-50 | 50-55 | 55-60 | >60 |
| 808 | 2440 | 2403 | 2154 | 2383 | 3059 |
| 3'UTR GC (%) | | | | | |
| <40 | 40-45 | 45-50 | 50-55 | 55-60 | >60 |
| 5569 | 2065 | 1745 | 1626 | 1346 | 865 |

**B**

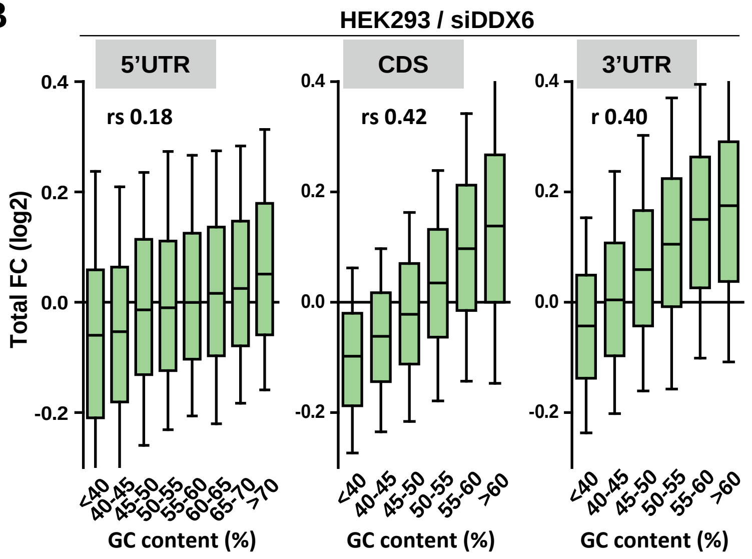


**C**

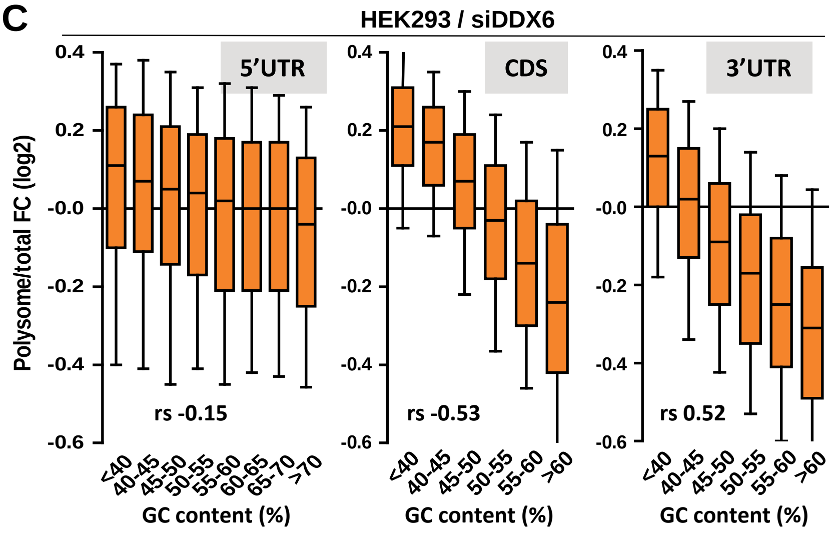

### Figure S6

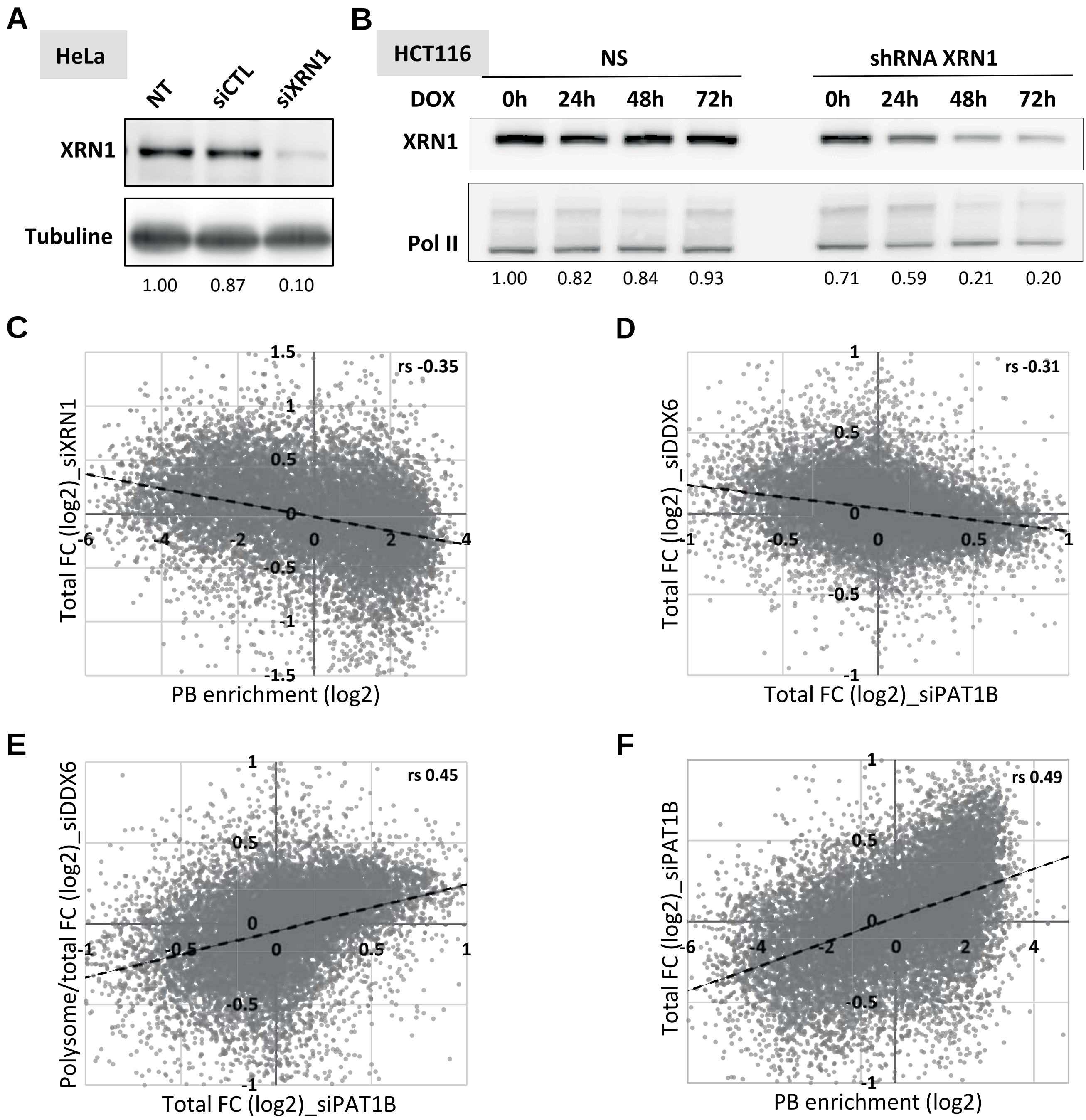

A
HeLa
NT
siCTL
siXRN1
XRN1
Tubuline
1.00
0.87
0.10
B
HCT116
NS
shRNA XRN1
DOX
0h
24h
48h
72h
0h
24h
48h
72h
XRN1
Pol II
1.00
0.82
0.84
0.93
0.71
0.59
0.21
0.20
C
rs -0.35
Total FC (log2)_siXRN1
PB enrichment (log2)
D
rs -0.31
Total FC (log2) _siDDX6
Total FC (log2)_siPAT1B
E
rs 0.45
Polysome/total FC (log2)_siDDX6
Total FC (log2)_siPAT1B
F
rs 0.49
Total FC (log2)_siPAT1B
PB enrichment (log2)

### Figure S7

**A**

| | Number of transcripts | | | | |
|---|---|---|---|---|---|
| | HEK293 siDDX6 | K562 shDDX6 | HeLa siXRN1 | HEK293 siPAT1B | HEK293 PBs |
| SMG6 | 220 | 217 | 211 | 211 | 211 |
| YTHDF2 | 1249 | 1182 | 1240 | 1283 | 1300 |
| G4 | 2330 | 1994 | 1886 | 2487 | 2362 |
| TOP | 104 | 102 | 102 | 104 | 104 |
| ARE | 2057 | 1680 | 1702 | 2140 | 2063 |
| HUR | 2664 | 2459 | 2521 | 2586 | 2601 |
| TTP | 1601 | 1489 | 1507 | 1535 | 1549 |
| FXR1 | 562 | 534 | 551 | 569 | 577 |
| FXR2 | 1413 | 1360 | 1390 | 1426 | 1449 |
| FMR1 | 658 | 637 | 648 | 673 | 682 |
| CPE | 2152 | 1757 | 1875 | 2208 | 2188 |
| PUM1 | 1207 | 1107 | 1126 | 1169 | 1158 |
| PUM2 | 657 | 603 | 632 | 662 | 669 |
| IGF2BP1 | 1330 | 1277 | 1312 | 1359 | 1371 |
| IGF2BP2 | 2449 | 2323 | 2387 | 2501 | 2528 |
| IGF2BP3 | 2736 | 2552 | 2657 | 2795 | 2823 |
| Mov10 | 2305 | 2159 | 2244 | 2344 | 2370 |
| 4E-T | 991 | 808 | 888 | 1005 | 992 |
| ATXN2 | 700 | 679 | 691 | 705 | 718 |

**B**

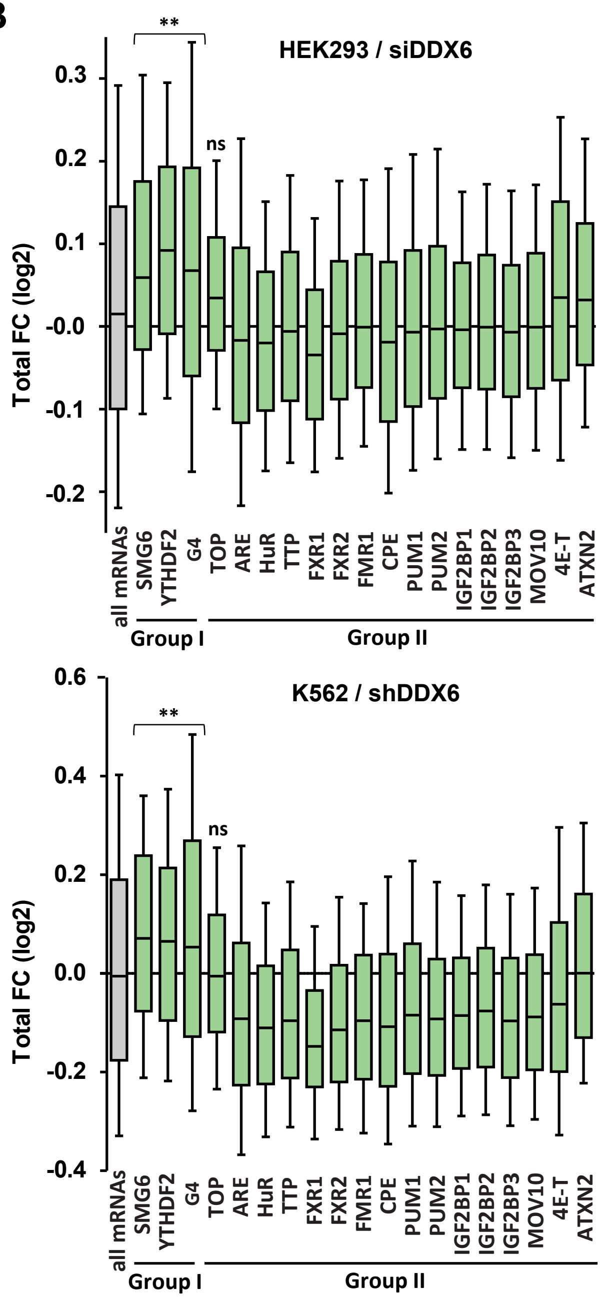


**C**

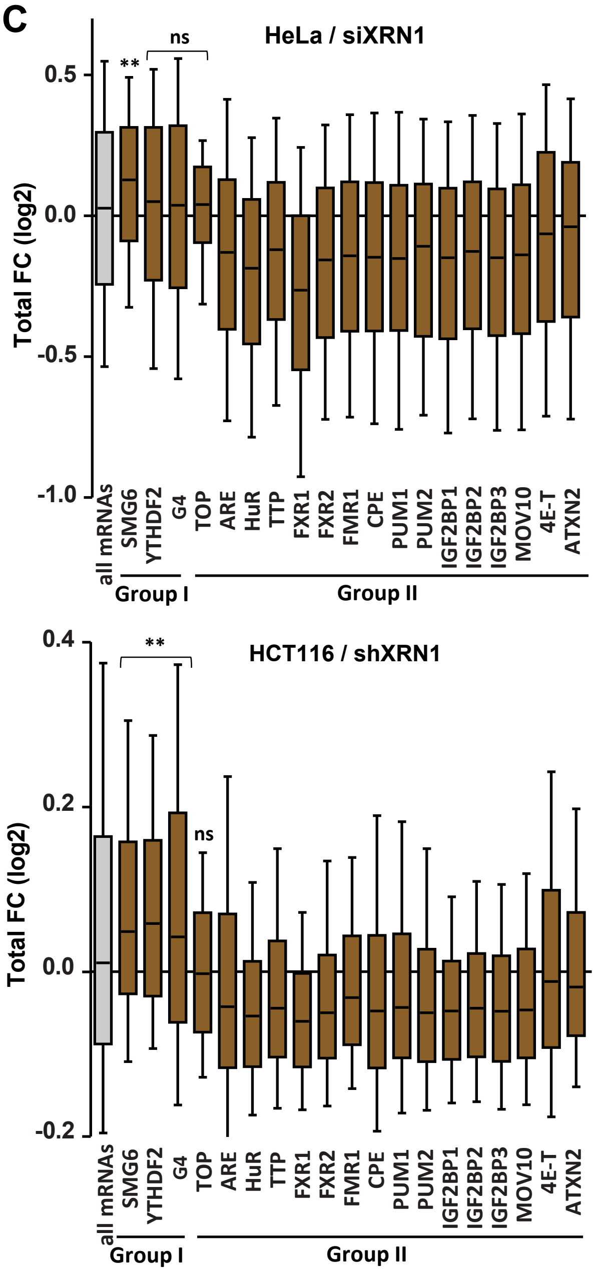


**D**

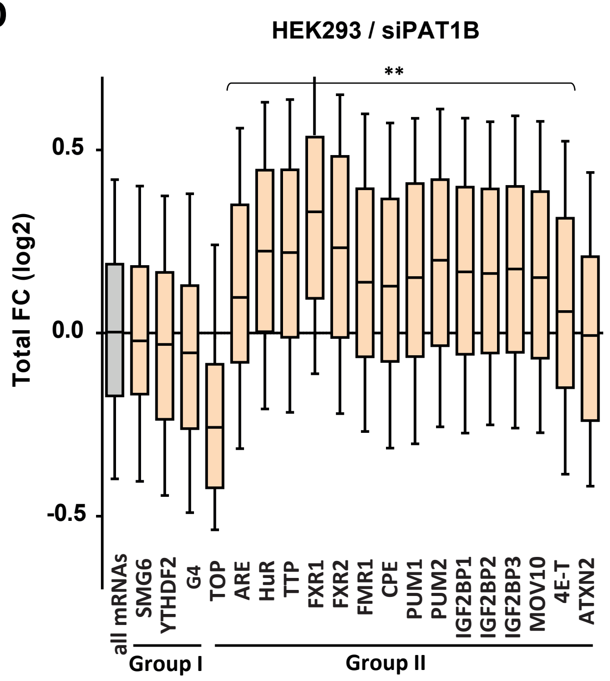

Figure S8

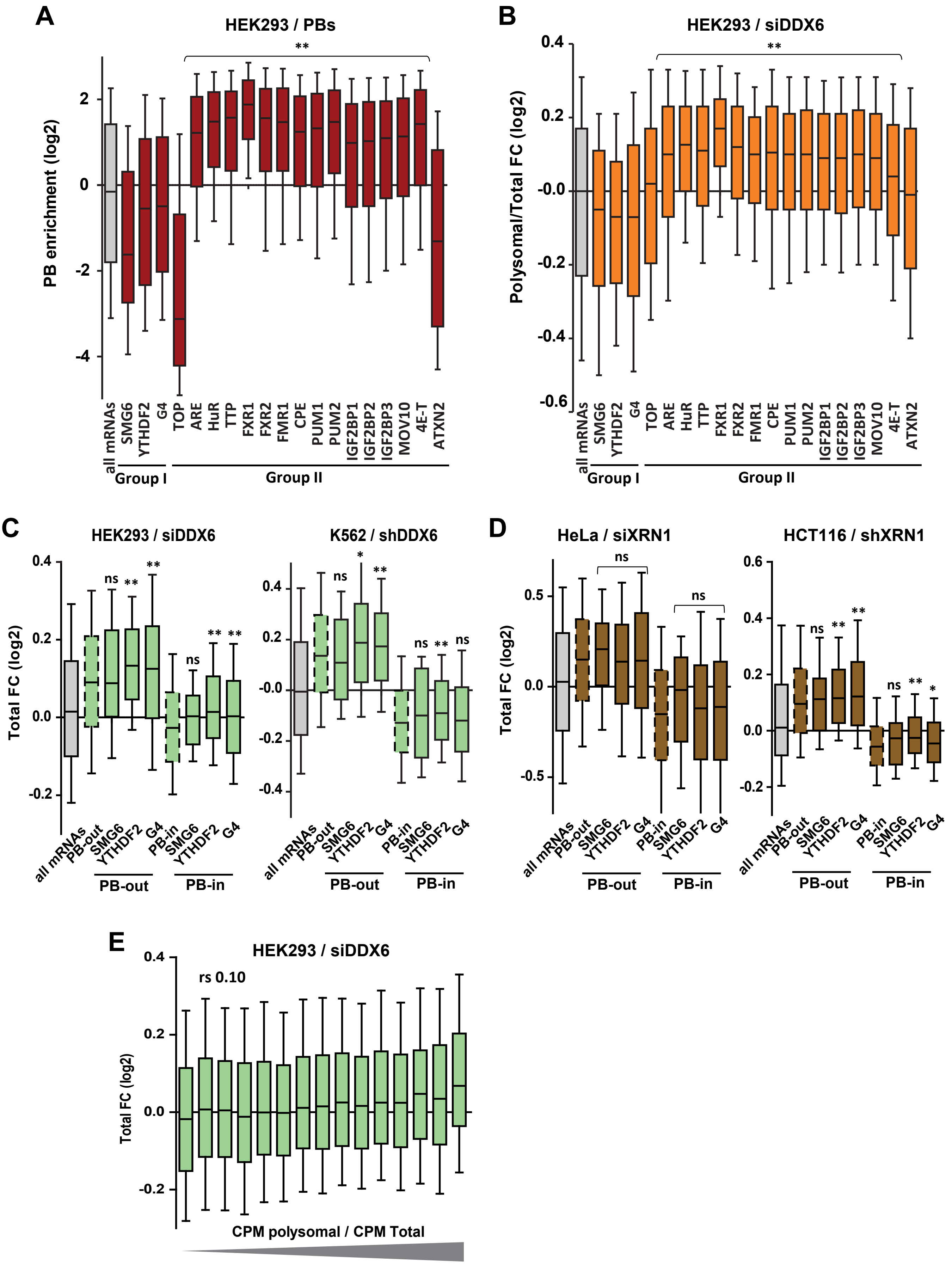
A
HEK293 / PBs
PB enrichment (log2)
2
0
-2
-4
**
all mRNAs
SMG6
YTHDF2
G4
TOP
ARE
HuR
TTP
FXR1
FXR2
FMR1
CPE
PUM1
PUM2
IGF2BP1
IGF2BP2
IGF2BP3
MOV10
4E-T
ATXN2
Group I
Group II
B
HEK293 / siDDX6
Polysomal/Total FC (log2)
0.4
0.2
-0.0
-0.2
-0.4
-0.6
**
all mRNAs
SMG6
YTHDF2
G4
TOP
ARE
HuR
TTP
FXR1
FXR2
FMR1
CPE
PUM1
PUM2
IGF2BP1
IGF2BP2
IGF2BP3
MOV10
4E-T
ATXN2
Group I
Group II
C
HEK293 / siDDX6
K562 / shDDX6
Total FC (log2)
0.4
0.2
0.0
-0.2
-0.0
-0.4
ns
**
*
all mRNAs
PB-out
SMG6
YTHDF2
G4
PB-in
SMG6
YTHDF2
G4
PB-out
PB-in
D
HeLa / siXRN1
HCT116 / shXRN1
Total FC (log2)
0.5
0.0
-0.5
0.6
0.4
0.2
-0.2
ns
**
*
all mRNAs
PB-out
SMG6
YTHDF2
G4
PB-in
SMG6
YTHDF2
G4
PB-out
PB-in
E
HEK293 / siDDX6
rs 0.10
Total FC (log2)
0.4
0.2
0.0
-0.2
CPM polysomal / CPM Total

#### Figure S9

**A**

| CLIP | Target number |
|---|---|
| AGO1 | 1043 |
| AGO2 | 421 |
| AGO3 | 1002 |
| AGO4 | 98 |

**B**

| miRNA | target number |
|---|---|
| hsa-miR-93-5p | 1127 |
| hsa-miR-19b-3p | 669 |
| hsa-miR-106b-5p | 997 |
| hsa-let-7a-5p | 238 |
| hsa-miR-19a-3p | 535 |
| hsa-miR-10a-5p | 432 |
| hsa-miR-17-5p | 1079 |
| hsa-miR-92a-3p | 1307 |
| hsa-miR-20a-5p | 985 |
| hsa-let-7f-5p | 351 |
| hsa-miR-21-5p | 561 |
| hsa-miR-30e-5p | 325 |
| hsa-miR-25-3p | 473 |
| hsa-miR-186-5p | 674 |
| hsa-miR-101-3p | 362 |
| hsa-miR-301a-3p | 369 |
| hsa-miR-18a-5p | 248 |
| hsa-miR-10b-5p | 296 |
| hsa-miR-16-5p | 1428 |
| hsa-miR-99a-5p | 126 |
| hsa-miR-99b-5p | 50 |
| hsa-miR-24-3p | 766 |

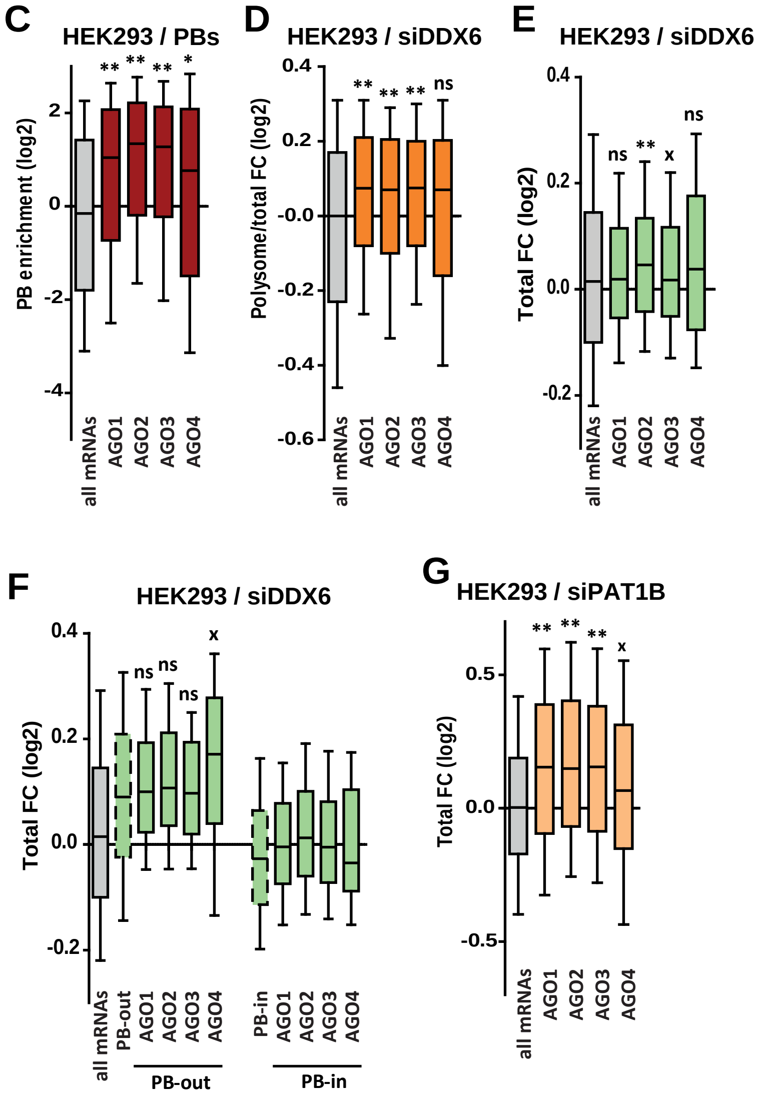


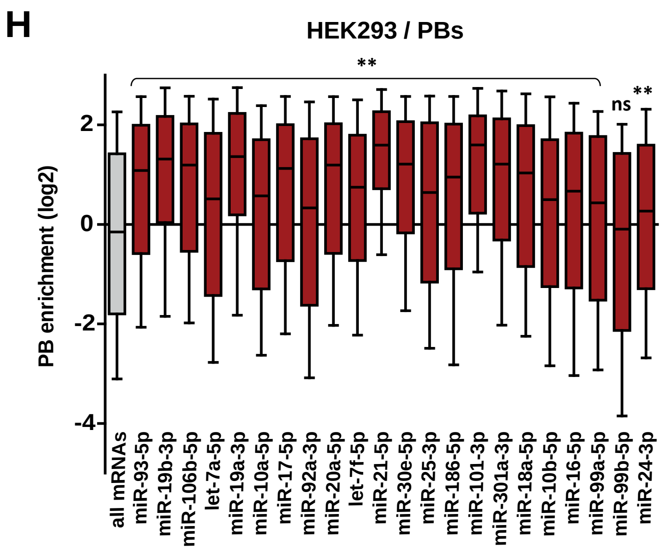

#### Figure S10

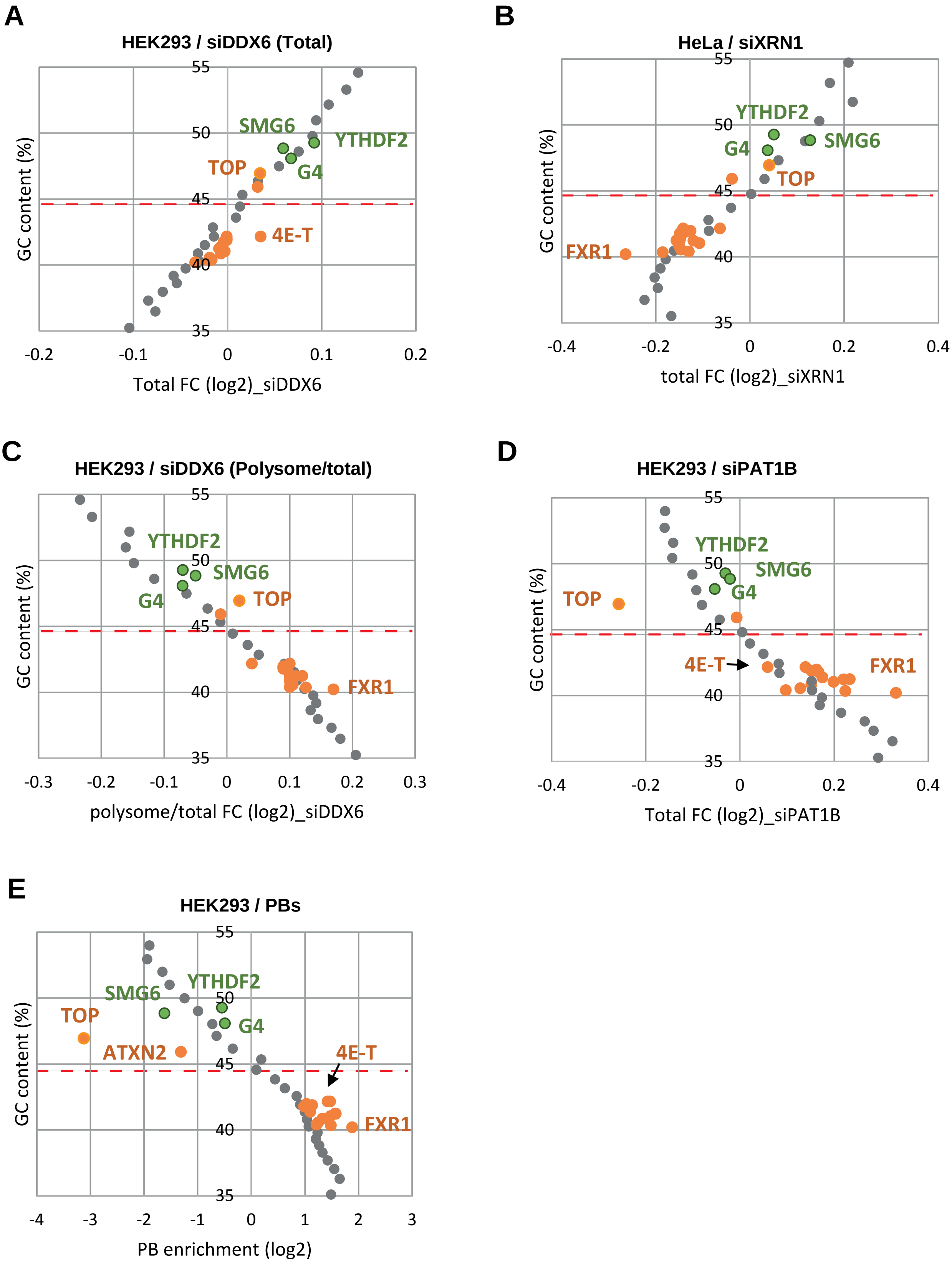

A
HEK293 / siDDX6 (Total)
GC content (%)
55
50
45
40
35
-0.2
-0.1
0
0.1
0.2
Total FC (log2)_siDDX6
SMG6
YTHDF2
TOP
G4
4E-T
B
HeLa / siXRN1
GC content (%)
55
50
45
40
35
-0.4
-0.2
0
0.2
0.4
total FC (log2)_siXRN1
YTHDF2
G4
SMG6
TOP
FXR1
C
HEK293 / siDDX6 (Polysome/total)
GC content (%)
55
50
45
40
35
-0.3
-0.2
-0.1
0
0.1
0.2
0.3
polysome/total FC (log2)_siDDX6
YTHDF2
SMG6
G4
TOP
FXR1
D
HEK293 / siPAT1B
GC content (%)
55
50
45
40
35
-0.4
-0.2
0
0.2
0.4
Total FC (log2)_siPAT1B
YTHDF2
SMG6
TOP
G4
4E-T
FXR1
E
HEK293 / PBs
GC content (%)
55
50
45
40
35
-4
-3
-2
-1
0
1
2
3
PB enrichment (log2)
YTHDF2
SMG6
TOP
G4
ATXN2
4E-T
FXR1

### Figure S11

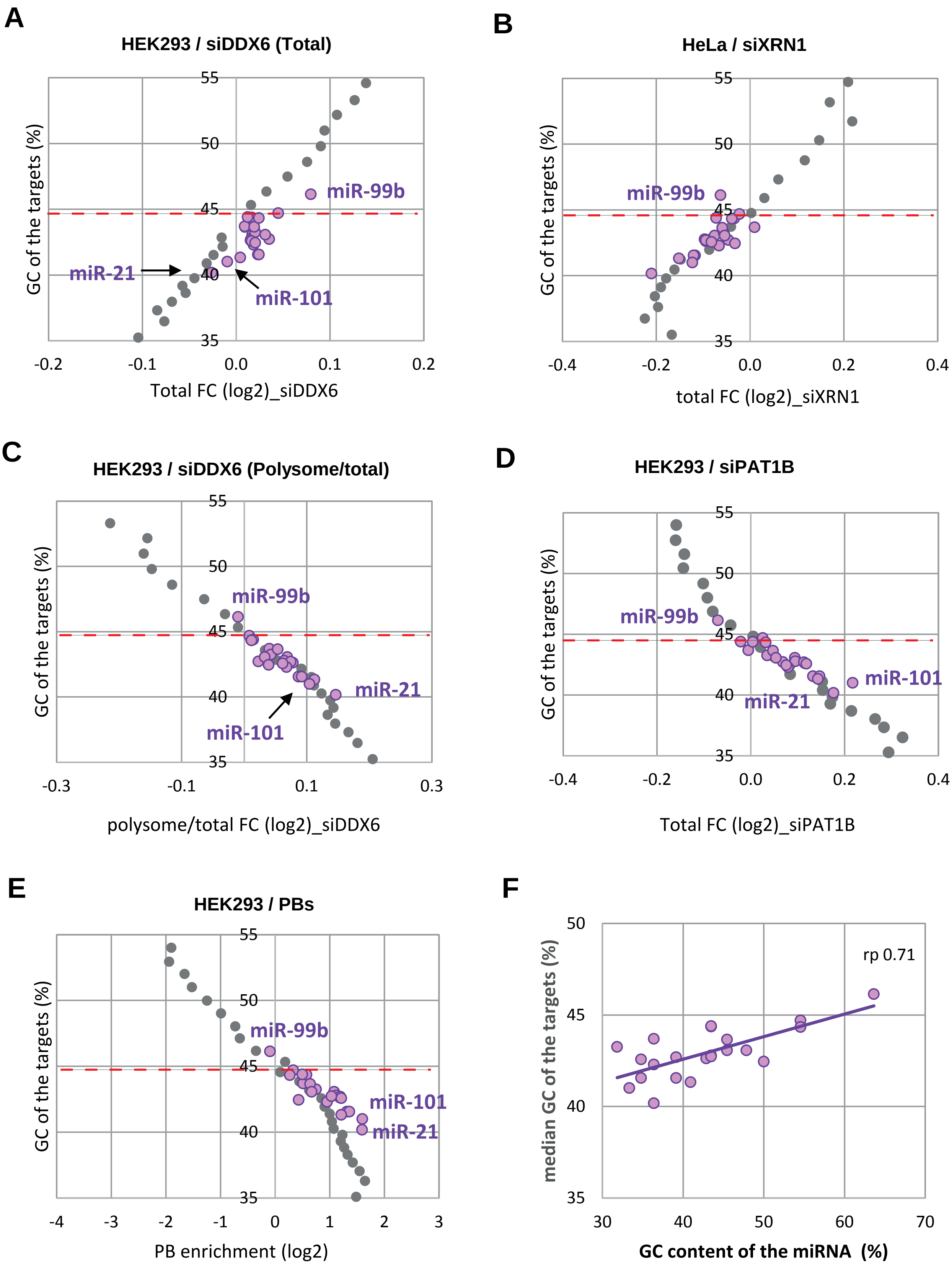
A
HEK293 / siDDX6 (Total)
GC of the targets (%)
55
50
45
40
35
-0.2
-0.1
0.0
0.1
0.2
Total FC (log2)_siDDX6
miR-99b
miR-21
miR-101
B
HeLa / siXRN1
GC of the targets (%)
55
50
45
40
35
-0.4
-0.2
0.0
0.2
0.4
total FC (log2)_siXRN1
miR-99b
C
HEK293 / siDDX6 (Polysome/total)
GC of the targets (%)
55
50
45
40
35
-0.3
-0.1
0.1
0.3
polysome/total FC (log2)_siDDX6
miR-99b
miR-21
miR-101
D
HEK293 / siPAT1B
GC of the targets (%)
55
50
45
40
35
-0.4
-0.2
0.0
0.2
0.4
Total FC (log2)_siPAT1B
miR-99b
miR-101
miR-21
E
HEK293 / PBs
GC of the targets (%)
55
50
45
40
35
-4
-3
-2
-1
0
1
2
3
PB enrichment (log2)
miR-99b
miR-101
miR-21
F
median GC of the targets (%)
50
45
40
35
30
40
50
60
70
GC content of the miRNA (%)
rp 0.71

**Figure S12**

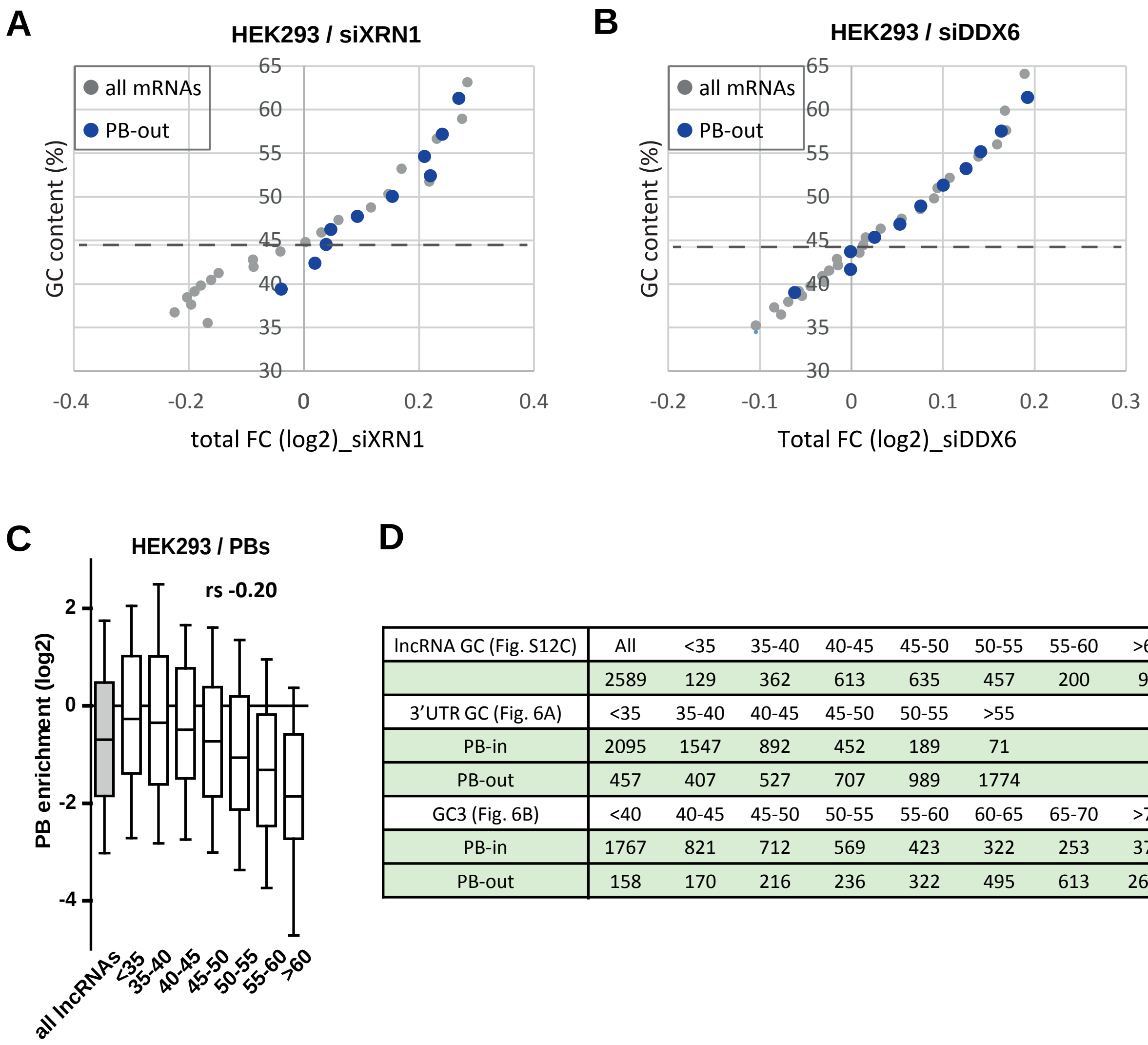


**D**

| lncRNA GC (Fig. S12C) | All | <35 | 35-40 | 40-45 | 45-50 | 50-55 | 55-60 | >60 |
|---|---|---|---|---|---|---|---|---|
| | 2589 | 129 | 362 | 613 | 635 | 457 | 200 | 95 |
| 3'UTR GC (Fig. 6A) | <35 | 35-40 | 40-45 | 45-50 | 50-55 | >55 | | |
| PB-in | 2095 | 1547 | 892 | 452 | 189 | 71 | | |
| PB-out | 457 | 407 | 527 | 707 | 989 | 1774 | | |
| GC3 (Fig. 6B) | <40 | 40-45 | 45-50 | 50-55 | 55-60 | 60-65 | 65-70 | >70 |
| PB-in | 1767 | 821 | 712 | 569 | 423 | 322 | 253 | 379 |
| PB-out | 158 | 170 | 216 | 236 | 322 | 495 | 613 | 2651 |

### Figure S13

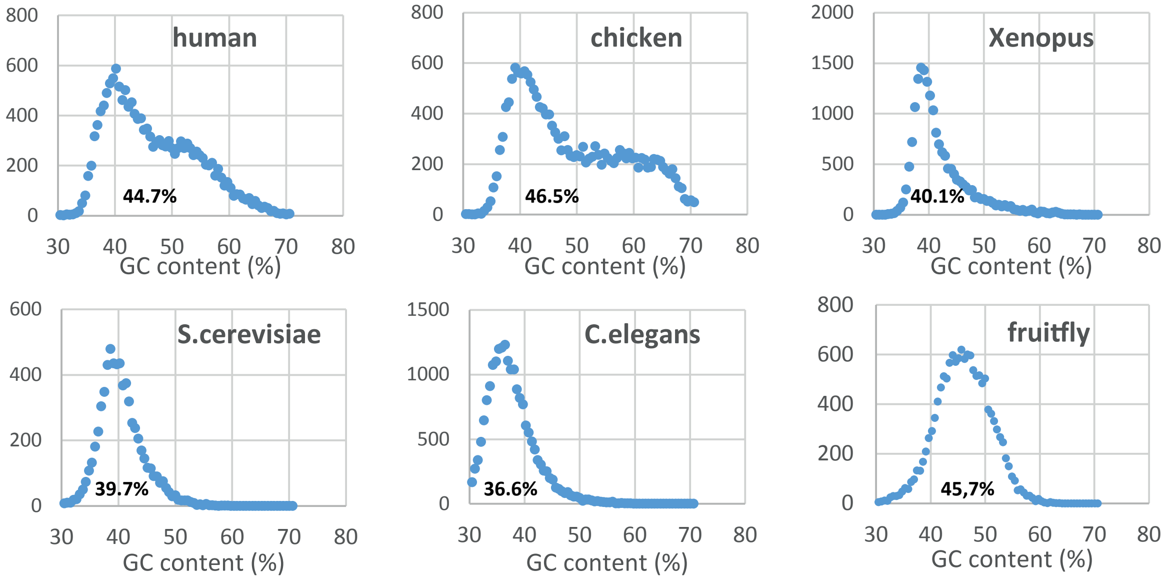
human
44.7%
chicken
46.5%
Xenopus
40.1%
S.cerevisiae
39.7%
C.elegans
36.6%
fruitfly
45,7%
GC content (%)
0
200
400
600
800
500
1000
1500
2000
30
40
50
60
70
80
